# Dissecting mutational mechanisms underpinning signatures caused by replication errors and endogenous DNA damage

**DOI:** 10.1101/2020.08.04.234245

**Authors:** Xueqing Zou, Gene Ching Chiek Koh, Arjun Scott Nanda, Andrea Degasperi, Katie Urgo, Theodoros I. Roumeliotis, Chukwuma A Agu, Lucy Side, Glen Brice, Vanesa Perez-Alonso, Daniel Rueda, Cherif Badja, Jamie Young, Celine Gomez, Wendy Bushell, Rebecca Harris, Jyoti S. Choudhary, Josef Jiricny, William C Skarnes, Serena Nik-Zainal

**Affiliations:** Academic Department of Medical Genetics, School of Clinical Medicine, University of Cambridge, Cambridge CB2 9NB, UK; MRC Cancer Unit, University of Cambridge, Cambridge CB2 0XZ, UK; Wellcome Trust Sanger Institute, Hinxton CB10 1SA, UK; The Institute of Cancer Research, Chester Beatty Laboratories, London SW3 6JB, UK; UCL Institute for Women’s Health, Great Ormond Street Hospital, London WC1N 3JH, UK; Wessex Clinical Genetics Service, Mailpoint 627, Princess Anne Hospital, Coxford Road, Southampton, SO16 5YA, UK; Southwest Thames Regional Genetics Service, St George’s University of London, Cranmer Terrace, London, SW17 ORE, UK; Pediatrics Department, Doce de Octubre University Hospital, i+12 Research Institute, Madrid, Spain; Hereditary Cancer Laboratory, Doce de Octubre University Hospital, i+12 Research Institute, Madrid, Spain; Institute of Molecular Life Sciences of the University of Zurich and Institute of Biochemistry of the ETH Zurich, Otto-Stern-Weg 3, Zurich 8093, Switzerland; The Jackson Laboratory for Genomic Medicine, 10 Discovery Drive, Farmington, Connecticut, 06032, USA

**Keywords:** DNA replicative/repair deficiency, Mutagenesis, Mutational signatures, Whole genome sequencing, CRISPR-Cas9

## Abstract

Mutational signatures are imprints of pathophysiological processes arising through tumorigenesis. Here, we generate isogenic CRISPR-Cas9 knockouts (Δ) of 43 genes in human induced pluripotent stem cells, culture them in the absence of added DNA damage, and perform wholegenome sequencing of 173 daughter subclones. Δ*OGG1*, Δ*UNG*, Δ*EXO1*, Δ*RNF168*, Δ*MLH1*, Δ*MSH2*, Δ*MSH6*, Δ*PMS1*, and Δ*PMS2* produce marked mutational signatures indicative of being critical mitigators of endogenous DNA changes. Detailed analyses reveal that 8-oxo-dG removal by different repair proteins is sequence-context-specific while uracil clearance is sequencecontext-independent. Signatures of mismatch repair (MMR) deficiency show components of C>A transversions due to oxidative damage, T>C and C>T transitions due to differential misincorporation by replicative polymerases, and T>A transversions for which we propose a ‘reverse template slippage’ model. Δ*MLH1*, Δ*MSH6*, and Δ*MSH2* signatures are similar to each other but distinct from Δ*PMS2*. We validate these gene-specificities in cells from patients with Constitutive Mismatch Repair Deficiency Syndrome. Based on these experimental insights, we develop a classifier, MMRDetect, for improved clinical detection of MMR-deficient tumors.

## Introduction

Somatic mutations arising through endogenous and exogenous processes mark the genome with distinctive patterns, termed mutational signatures^1–4^. While there have been advancements in analytical aspects of deriving mutational signatures from human cancers^5–7^, there is an emerging need for experimental substantiation, elucidating etiologies and mechanisms underpinning these mutational patterns^8–11^. Cellular models have been used to systematically study mutagenesis arising from exogenous sources of DNA damage^8,11^. Next, it is essential to experimentally explore genome-wide mutagenic consequences of endogenous sources of DNA damage, in the absence of external DNA damaging agents.

Lindahl noted that water and oxygen, essential molecules for living organisms, are some of the most mutagenic elements to DNA^12^. His seminal work demonstrated that spontaneous DNA lesions occur through endogenous biochemical activities such as hydrolysis and oxidation. Errors at replication are also an enormous potential source of DNA changes. Fortuitously, our cells are equipped with DNA repair pathways that constantly mitigate this endogenous damage^13,14^. In this work, we combine CRISPR-Cas9-based biallelic knockouts of a selection of DNA replicative/repair genes in human induced Pluripotent Stem Cells (hiPSCs), whole-genome sequencing(WGS), and in-depth analysis of experimentally-generated data, to obtain mechanistic insights into mutation formation. It is beyond the scope of this manuscript to study all DNA repair genes. Thus, we have focused on 42 DNA repair gene knockouts successfully generated through semi-high-throughput methods. We also perform comparisons between experimental data and previously generated cancer-derived signatures.

While there is enormous literature regarding DNA repair pathways and complex protein interactions that are involved in maintaining genomic integrity^15–20^, here we focus on directly mapping whole-genome mutational outcomes associated with human DNA repair defects, critically, in the absence of any applied, external damage. This study, therefore, allows us to identify replicative/repair genes that are fundamentally important to genome maintenance against endogenous sources of DNA damage.

## Results

### Biallelic knockouts of DNA repair genes

We knocked out (Δ) 42 genes involved in DNA repair/replicative pathways and an unrelated control gene, *ATP2B4* (Figures 1A and 1B, Table S1). Two knockout genotypes were generated per gene except for *EXO1, MSH2, TDG, MDC1*, and *REV1*, where only one knockout genotype was obtained. All parental knockout lines analyzed below were grown over 15 days under normoxic conditions (~20% oxygen). For each genotype, two single-cell subclones were derived for whole-genome sequencing (WGS), aiming for four sequenced subclones per edited gene (Figure 1A). For single genotype genes, three subclones were derived for Δ*EXO1* and Δ*MSH2*, and four for Δ*TDG*, Δ*MDC1*, and Δ*REV1*.

**Figure 1.**
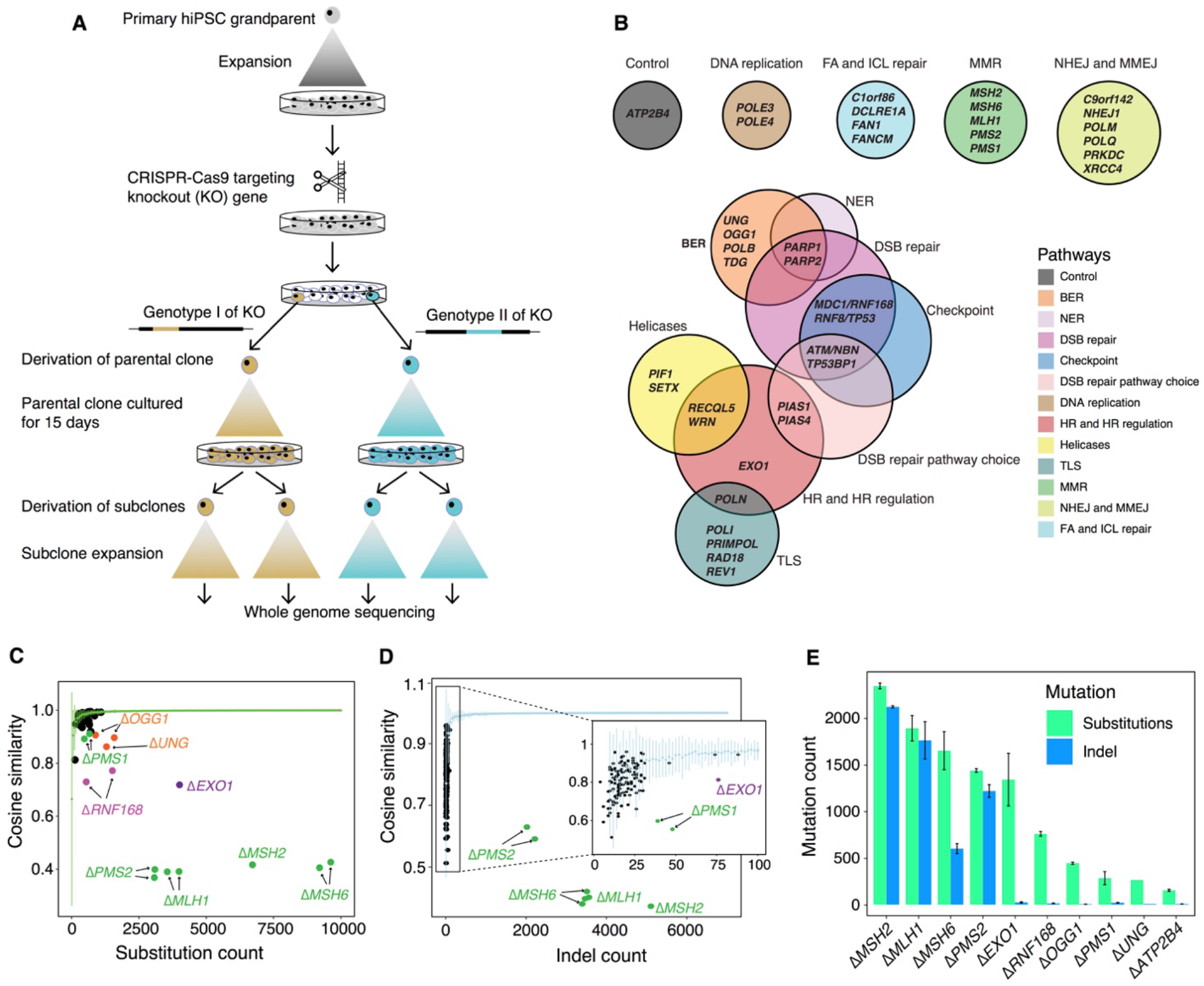
Mutational consequences of DNA repair gene knockouts. (A) Experimental workflow from isolation of gene knockouts to generating subclones for WGS. (B) Forty-three genes were knocked out, including 42 DNA repair/replication genes and one control gene (*ATP2B4*). (C) Distinguishing substitution profiles of control subclones and knockout subclones. Green line shows the cosine similarities between bootstrapped profiles of controls against aggregated control substitution profile. (D) Distinguishing indel profile of control subclones and knockout subclones. Light blue line shows the cosine similarities between bootstrapped indel profiles of controls against aggregated control indel profile. (E) De novo mutation number of knockout subclones cultured for 15 days. Bars and error bars represent mean ± SD of subclone observations.

A total of 173 subclones were obtained from 78 genotyped knockouts of 43 genes (Table S2). All subclones were sequenced to an average depth of ~25-fold. Short-read sequences were aligned to human reference genome assembly GRCh37/hg19. All classes of somatic mutations were called, subtracting variation of the primary hiPSC parental clone (Table S2-S3, Figure S1–S2, pilot results in Supplementary Note 1). Rearrangements were too infrequent to decipher specific patterns.

We confirmed that mutational outcomes were neither due to off-target edits nor to the acquisition of new driver mutations (Online Methods). We verified that knockouts were biallelic, confirmed this further by protein mass spectrometry, and ensured that subclones were derived from single cells in all comparative analyses (Online Methods).

### Mutational consequences of gene knockouts

We reasoned that under these controlled experimental settings, if simply knocking-out a gene (in the absence of providing additional DNA damage) could produce a signature, then the gene is critical to maintaining genome stability from endogenous sources of DNA damage. It would manifest an increased mutation burden above background and/or an altered mutation profile (Figure S3). We found background substitution and indel mutagenesis associated with growing cells in culture occurred at ~150 substitutions and ~10 indels per genome and was comparable across all subclones.

To address potential uncertainty associated with the relatively small number of subclones per knockout and variable mutation counts in each gene knockout (Methods), we generated bootstrapped control samples with variable mutation burdens (50-10,000). We calculated cosine similarities between each bootstrapped sample and the background control (Δ*ATP2B4*) mutational signature (mean and standard deviations). A cosine similarity close to 1.0 indicates that the mutation profile of the bootstrapped sample is near-identical to the control signature. Cosine similarities could thus be considered across a range of mutation burdens (green line in Figure 1C and light blue line in Figure 1D). We next calculated cosine similarities between knockout profiles and controls (colored dots in Figures 1C and 1D). A knockout experiment that does not fall within the expected distribution of cosine similarities implies a mutation profile distinct from controls, i.e., the gene knockout is associated with a signature. For substitution signatures, two additional dimensionality reduction techniques, namely, contrastive principal component analysis (cPCA)^21^ and t-Distributed Stochastic Neighbor Embedding (t-SNE)^22^ were also applied to secure high confidence mutational signatures (Figure S4, Methods). This stringent series of steps would likely dismiss weaker signals and thus be highly conservative at calling mutational signatures. These conservative methods were also applied to identify indel signatures (Methods).

We identified nine single substitution, two double substitution and six indel signatures. Three gene knockouts, Δ*OGG1*, Δ*UNG,* and Δ*RNF168*, produced only substitution signatures. Six gene knockouts, Δ*MSH2*, Δ*MSH6*, Δ*MLH1*, Δ*PMS2*, Δ*EXO1*, and Δ*PMS1*, presented substitution and indel signatures. Δ*EXO1* and Δ*RNF168* also produced double substitution patterns. The average *de novo* mutation burden accumulated for these nine knockouts (Figure 1E) ranged between 250-2,500 for substitutions and 5-2,100 for indels. Based on cell proliferation assays, mutation rates for each knockout were calculated and ranged between 6-129 substitutions and 0.39-126 indels per cell division (Table S4). In the following sections, we dissect these experimentally-generated mutational signatures. We compare them to one another and to previously published human cancer-derived mutational signatures, to gain insights into the sources of endogenous DNA damage and mutational mechanisms.

### Safeguarding the genome from oxidative DNA damage

Oxygen can generate reactive oxygen species (ROS) and oxidative DNA lesions. The commonest is 8-oxo-2’-deoxyguanosine (8-oxo-dG), although over 25 oxidative DNA lesions are known^23^. 8-oxo-dG is predominantly repaired by Base Excision Repair (BER). A pervasive mutational signature observed in cell-based experiments has been speculated as due to culture-related oxidative damage^9,11^. It is similar to a mutational signature identified in adrenocortical cancers and neuroblastomas, called RefSig18^24^ or SBS18^6^. Biallelic loss of MutY DNA-glycosylase (MUTYH), which excises adenines inappropriately paired with 8-oxo-dG, has also been reported to generate a hypermutated version of a similar signature^25^. It is unclear whether other genes responsible for removing oxidative damage would also result in these characteristic patterns.

8-Oxoguanine glycosylase (OGG1) is responsible for the excision of 8-oxo-dG^26^. Thus, an Δ*OGG1* signature would be an undisputed pattern of 8-oxo-dG-related damage. Δ*OGG1* produced a marked G>T/C>A pattern particularly at T<u>G</u>C>T<u>T</u>C/G<u>C</u>A>G<u>A</u>A with additional peaks at T<u>G</u>T>T<u>T</u>T/A<u>C</u>A>A<u>A</u>A, C<u>G</u>A>C<u>T</u>A/G<u>C</u>T>G<u>A</u>T and A<u>G</u>A>A<u>T</u>A/T<u>C</u>T>T<u>A</u>T (Figure 2A), similar to the culture-related signature and RefSig18/SBS18 (Figure 2A and 2B). Not only does this support the hypothesis that those signatures are due to oxidative damage, it specifically implicates 8-oxo-dG. We further expanded signature channels by considering ±2 bases at the 5’ and 3’ positions around the mutated base. Higher-resolution assessment of the most dominant peak at T<u>G</u>C>T<u>T</u>C/G<u>C</u>A>G<u>A</u>A in Δ*OGG1* showed an almost identical pattern to control samples carrying culture-related signatures and SBS18 (cosine similarity (cossim): > 0.9, Figure 2C, S5A), strengthening the argument that the G>T/C>A transversions observed in cultured cells and SBS18 are indeed mainly caused by 8-oxo-dG-related damage.

**Figure 2.**
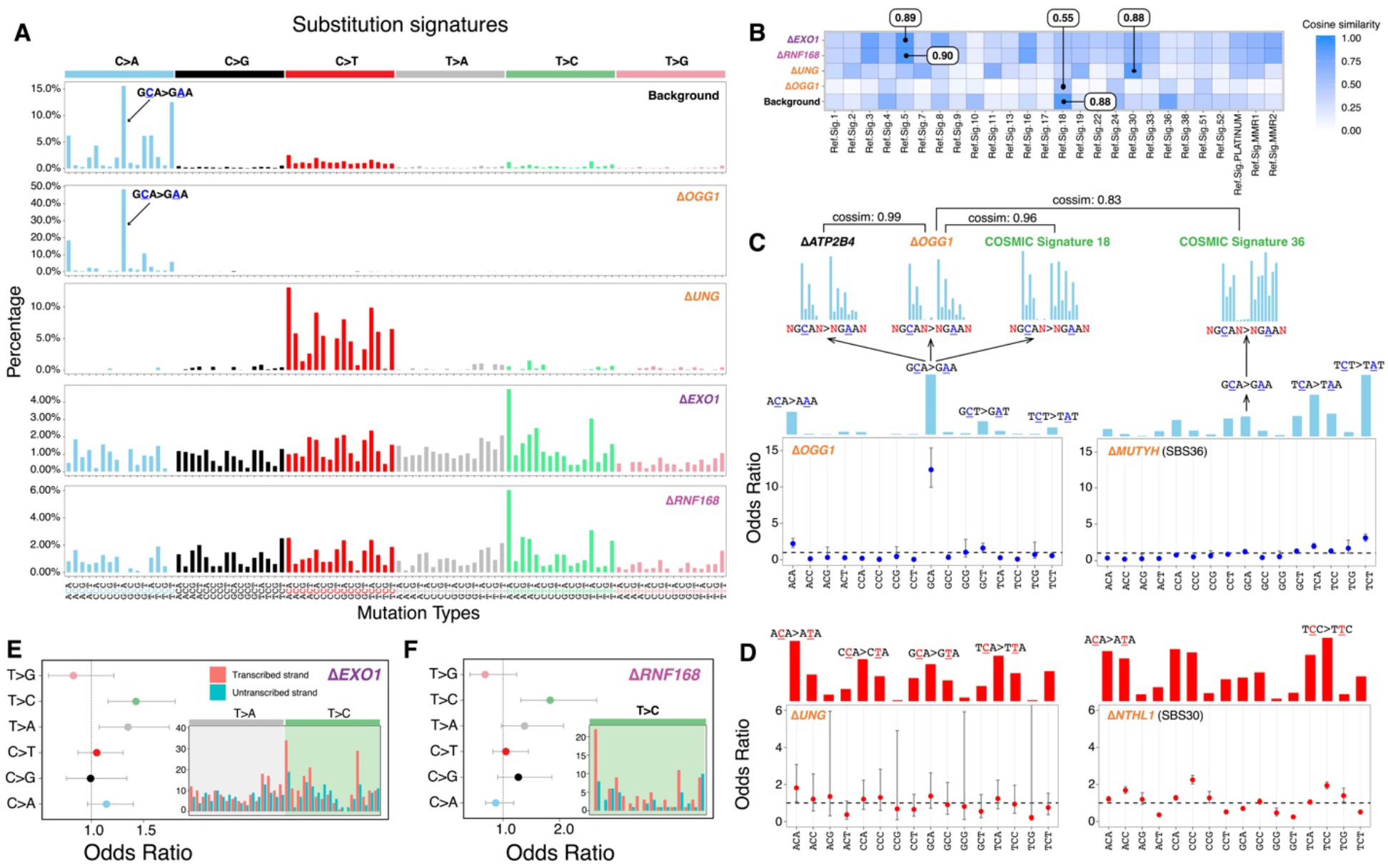
Safeguarding the genome from oxidative damage and cytosine deamination. (A) Substitution signatures of background mutagenesis (from control), Δ*OGG1*, Δ*UNG*, Δ*EXO1* and Δ*RNF168*. (B) Cosine similarity between mutational signature of gene knockouts and Cancer-derived mutational signatures^24^. (C) Odds ratio of C>A occurring at 16 trinucleotides for Δ*OGG1* and Δ*MUTYH* (SBS36)^6^. Calculation was corrected for distribution of trinucleotides in the reference genome. Odds ratio less than 1 with 95% confidence interval (CI) < 1 implies that C>A mutations at that particular trinucleotide are less likely to occur. The mutational profiles of C>A at GCA with ±2 flanking bases are shown for Δ*ATP2B4*, Δ*OGG1*, SBS18 and SBS36. (D) Odds ratio of C>T occurring at all 16 trinucleotides for Δ*UNG* and Δ*NTHL1* (SBS30)^6^. Transcriptional strand asymmetry of (E) Δ*EXO1* signature and (F) Δ*RNF168* signature. The insets show the count number of T>C/A>G mutations on transcribed and non-transcribed strands.

Δ*OGG1* signature is qualitatively analogous to the signature of Δ*MUTYH-related* adrenocortical cancers^25^ (recently renamed SBS36^6^), although the latter demonstrates hypermutator phenotypes and has its tallest peak at T<u>C</u>T instead (Figure 2C). These similarities are explained by related but distinct roles played by *OGG1* and *MUTYH* in repairing oxidation-related lesions: 8-oxo-dG can pair either with C or with A during DNA synthesis. 8-oxo-G/C mismatches are, however, not mutagenic and oxidized guanines are simply excised by OGG1 glycosylase^27^. By contrast, 8-oxo-G/A mismatches are first repaired by MutY-glycosylase, which removes the A and repair synthesis by pol-β or −λ inserts a C opposite the oxidized base. The resulting 8-oxo-G/C pair is then excised by OGG1 as outlined earlier. This mechanistic relatedness likely explains why mutational signatures of Δ*OGG1* and Δ*MUTYH* are qualitatively alike, if quantitatively dissimilar. Notably, that simple knockouts of *OGG1* or *MUTYH* can result in overt mutational phenotypes suggests that these genes are indispensable for maintaining the genome against endogenous oxidative damage.

Lastly, we examined Δ*OGG1* G>T/C>A mutations correcting for frequencies of the 16 trinucleotides in the reference genome and found that Δ*OGG1* is depleted of mutations at GG/CC dinucleotides (Figure 2C). Yet, prior literature reports 5’-G in GG and the first two Gs in GGG are more likely to be oxidized through intraduplex electron transfer reactions^28,29^. Therefore, one would expect to observe elevated G>T/C>A mutation burdens in GG-rich regions when key repair proteins such as OGG1 are defective. Our results may be explained by previous experiments which demonstrate that 8-oxo-dG excision rates by OGG1 are sequence-context dependent^30^: 8-oxo-dG excision at consecutive GG sequences is reported as inefficient compared to 5’-C<u>G</u>C/5’-G<u>C</u>G and 5’-A<u>G</u>C/5’-G<u>C</u>T because OGG1 employs a bend-and-flip strategy to recognize 8-oxoG^31–33^. Stacked adjacent 8-oxo-dGs have an increased kinetic barrier, preventing flipping out and removal of 8-oxo-dG^30^. While this may explain why OGG1 cannot repair oxidized guanines at GG/CC motifs, it remains unclear how these motifs are repaired as guanine oxidation does occur at such sites. At some GG/CC motifs, we suggest a possibility in the section on mismatch repair genes later.

### Maintaining cytosines from deamination to uracil

Deamination involves hydrolytic loss of an amine group. At CpG dinucleotides, deamination of 5-methylcytosine into thymine is a well-studied, universal process^34,35,36^. In human cancers, C>T mutagenesis at CpGs (Signature 1) manifests in many tumor types. Hypermutator phenotypes of C>T mutations at CpGs, however, have been reported in cancers with biallelic loss of methyl-binding domain 4 (*MBD4*)^37^. This example underscores a mutational process that is customarily under tight *MBD4* regulation, wherein its knockout uncovers the potential magnitude of unrepaired endogenous deamination.

Spontaneous hydrolytic cytosine deamination to uracil occurs more slowly at ~10^2^-10^3^/cell/day, catalyzed by cytidine deaminases, APOBECs^38^. Cytosine deamination to uracil is rectified by *UNG* (uracil-N-glycosylase) via BER^39^. Uracils that are not removed prior to replication can result in C>T mutations (Figure 2A). There are mutational signatures associated with enhanced enzymatic activity of APOBEC in many human cancers (Signatures 2 and 13). However, the consequence of *UNG* dysfunction is less clear. The Δ*UNG* signature comprised C>T transitions but was not focused at specific trinucleotide sequence contexts. When corrected for trinucleotide frequencies in the reference genome (Figure 2D), there was no preference observed across all trinucleotides, supporting a general role for UNG activity on all uracils regardless of sequence context. Δ*UNG* signature shows the greatest similarity to RefSig 30 (cossim 0.88) but is unlikely to be its source given that Signature 30 was previously associated with Δ*NTHL1*^40^. Both UNG and NTHL1 are BER glycosylases that process aberrant pyrimidines, which may explain similarities between these signatures. However, when corrected for trinucleotide frequencies in the reference genome, Δ*NTHL1* shows slight preference for ACC, CCC, and TCC trinucleotides in contrast to Δ*UNG.*

### Preserving thymines and adenines from T>C/A>G transitions

Two genes *EXO1* and *RNF168* with wide-ranging roles in repair/checkpoint pathways^41,42,43^ showed mutational signatures. *EXO1* encodes a 5’ to 3’ exonuclease with RNase H activity. Δ*EXO1* generated substitution, double-substitution, and indel signatures in hiPSC (Figures 2A and S6), consistent with a previous study of Δ*EXO1* in HAP1 lines^9^. In HAP1 cells, Δ*EXO1* had stronger C>A components, probably reflecting the difference in model systems. Δ*EXO1* also produced a double substitution pattern defined mainly by TC>AT, TC>AA and GC>AA mutations, and an indel signature characterized by 1 bp A/T insertions at long poly[d(A-T)] (>= 5 bp) and 1 bp deletions at short poly[d(A-T)] or poly[d(C-G)] (< 5 bp) (Figure S6).

*RNF168* encodes an E3 ubiquitin ligase involved in DNA double-strand break (DSB) repair^43^. It regulates 53BP1 recruitment to DNA double-strand breaks (DSBs) through ubiquitin-dependent signaling^44^. The substitution signature of Δ*RNF168* has two T>C peaks at A<u>T</u>A>A<u>C</u>A and T<u>T</u>A>T<u>C</u>A (Figure 2A) and shares similarity with Δ*EXO1* (cossim: 0.94). Double substitution patterns were defined by TC>AA and GC>AA mutations. Indel signatures were not seen for Δ*RNF168.*

Δ*EXO1* and Δ*RNF168* signatures are most similar to RefSig5 of cancer-derived signatures (Figure 2B and S7, cossim: 0.89-0.9), defined mainly by T>C/A>G substitutions. Additionally, Δ*EXO1* and Δ*RNF168* signatures show transcriptional strand bias for T>C/A>G mutations (Figure 2E and 2F), in particular, at A<u>T</u>A and T<u>T</u>A context, with bias for T>C on the transcribed strand (A>G on nontranscribed strand). This is also in-keeping with Signature 5, which exhibits transcriptional strand bias for T>C/A>G mutations. The etiology of Signature 5 is currently unknown, although a hypermutator phenotype has been reported in association with loss of *ERCC2^7^*. Due to its similarity to Δ*EXO1* and Δ*RNF168* signatures, the wide-ranging roles played by these proteins and the transcriptional strand bias observed, we speculate that Signature 5 has a complex origin, and may be associated with endogenous sources of DNA damage that are repaired by multiple proteins and repair pathways.

### Multiple endogenous sources of DNA damage managed by mismatch repair

Knockouts of five genes involved in the mismatch repair (MMR) pathway^45–47^, *MSH2, MSH6, MLH1*, *PMS2,* and *PMS1,* produced substitution and indel signatures (Figure 3A and 3B) but not double substitution signatures despite a previously reported association^6^. Δ*MLH1*, Δ*MSH2*, and Δ*MSH6* produced identical qualitative substitution signatures (cossim: 0.99) characterized by a single strong peak at C<u>C</u>T>C<u>A</u>T/A<u>G</u>G>A<u>T</u>G, and multiple peaks of C>T and T>C (Figure 3A). In contrast, Δ*PMS2* generated a signature of predominantly T>C transitions with a slight predominance at A<u>T</u>A, A<u>T</u>G, and C<u>T</u>G (Figure 3C). The single peak at C<u>C</u>T>C<u>A</u>T/A<u>G</u>G>A<u>T</u>G remains visible in the Δ*PMS2* substitution signature, albeit markedly reduced (10% to 3%). In addition, Δ*MSH2, ΔMSH6,* and Δ*MLH1* generated indel signatures dominated by A/T deletions at long repetitive sequences. In contrast, Δ*PMS2* produced similar amounts of A/T insertions and A/T deletions at long repetitive sequences (Figures 3B, S8 and S9). Δ*PMS1* generated A/T deletions only at long poly[d(A-T)] (>=5 bp) and long deletions (> 1bp) at repetitive sequences (Figure S9).

**Figure 3.**
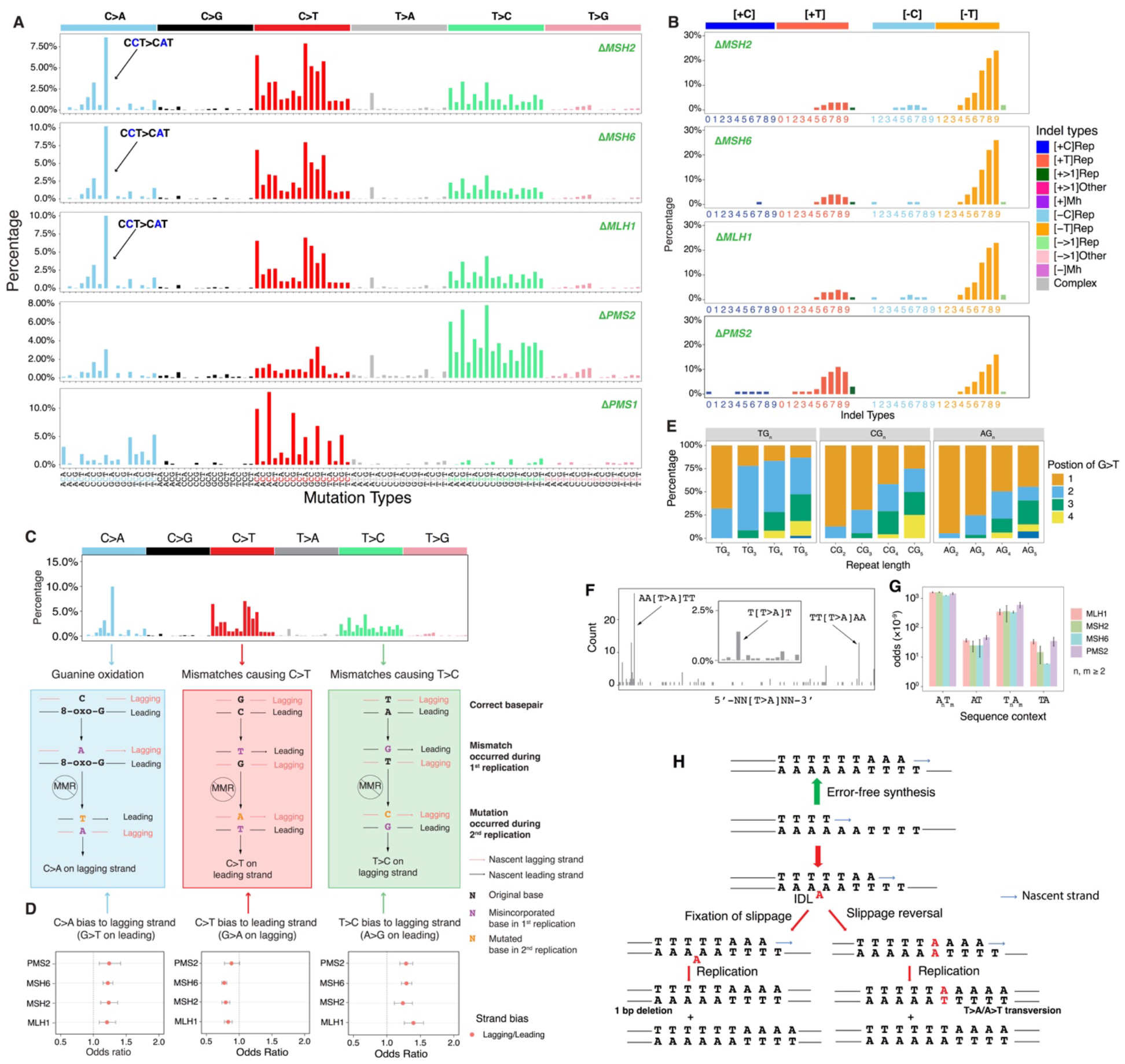
Multiple endogenous sources of DNA damage managed by mismatch repair. (A) Substitution and (B) indel signatures for five mismatch repair gene knockouts. The indel signature of ΔPMS1 is shown in Figure S9. (C) Dissection of DNA mismatch repair mutational signatures: C>A mutations believed to be due to unrepaired oxidative damage of guanine, and proposed mechanism of how DNA polymerase errors cause mis-incorporated bases that result in C>T and T>C. All other mismatch possibilities and their outcomes are demonstrated in Figure S10 The red and black strands represent lagging and leading strands, respectively. The arrowed strand is the nascent strand. (D) Replicative strand asymmetry observed for mutational signatures generated by four MMR gene knockouts. Data are represented as calculated odds ratio with 95% confidence interval. (E) The relative frequency of occurrence of G>T/C>A in polyG tracts for Δ*MSH6.* The count and relative frequency of occurrence of G>T/C>A in polyG tracts for Δ*MSH2* and Δ*MLH1* are shown in Figure S12. (F) T>A mutation frequency is highest at junctions of poly(A)poly(T) or poly(T)poly(A). (G) Odds for T>A mutations to occur at poly(A)poly(T) or poly(T)poly(A) are higher than AT sequences flanked by other nucleotides, corrected for sequence context through whole genome. Data are represented as mean ± SEM. (H) Putative models of T>A substitutions at poly(A)poly(T) or poly(T)poly(A) junctions due to template strand slippage and slippage reversal.

In-depth analysis of these mutational signatures allowed us to determine putative sources of endogenous DNA damage (Figure 3C) acted upon by MMR.

First, we consistently observed replication strand bias across Δ*MLH1*, Δ*MSH2*, Δ*MSH6*, and Δ*PMS2*: C>A on the lagging strand (equivalent to G>T leading strand bias), C>T on the leading strand (or G>A lagging) and T>C lagging (or A>G leading) (Figure 3D). Under our experimental settings where exogenous DNA damage was not administered, mismatches may be generated by DNA polymerases α, δ or ε during replication. In the absence of MMR, these lesions become permanently etched as mutations. To understand which replicative polymerases could be causing these mutations, we analyzed putative progeny of all twelve possible base/base mismatches (Figure S10). T/G mismatches are the most thermodynamically stable and represents the most frequent polymerase error^48^. Our assessment suggests that the predominance of T>C transitions on the lagging-strand can only be explained by misincorporation of T by lagging strand polymerases, pol-*a* and/or pol-*δ* leading to G/T mismatches (Figures 4C). Similarly, the observed bias for C>T transitions on the leading strand is likely to be predominantly caused by misincorporation of G on lagging strand by pol-*a* and/or pol-*δ* resulting in T/G mismatches (Figures 4C).

**Figure 4.**
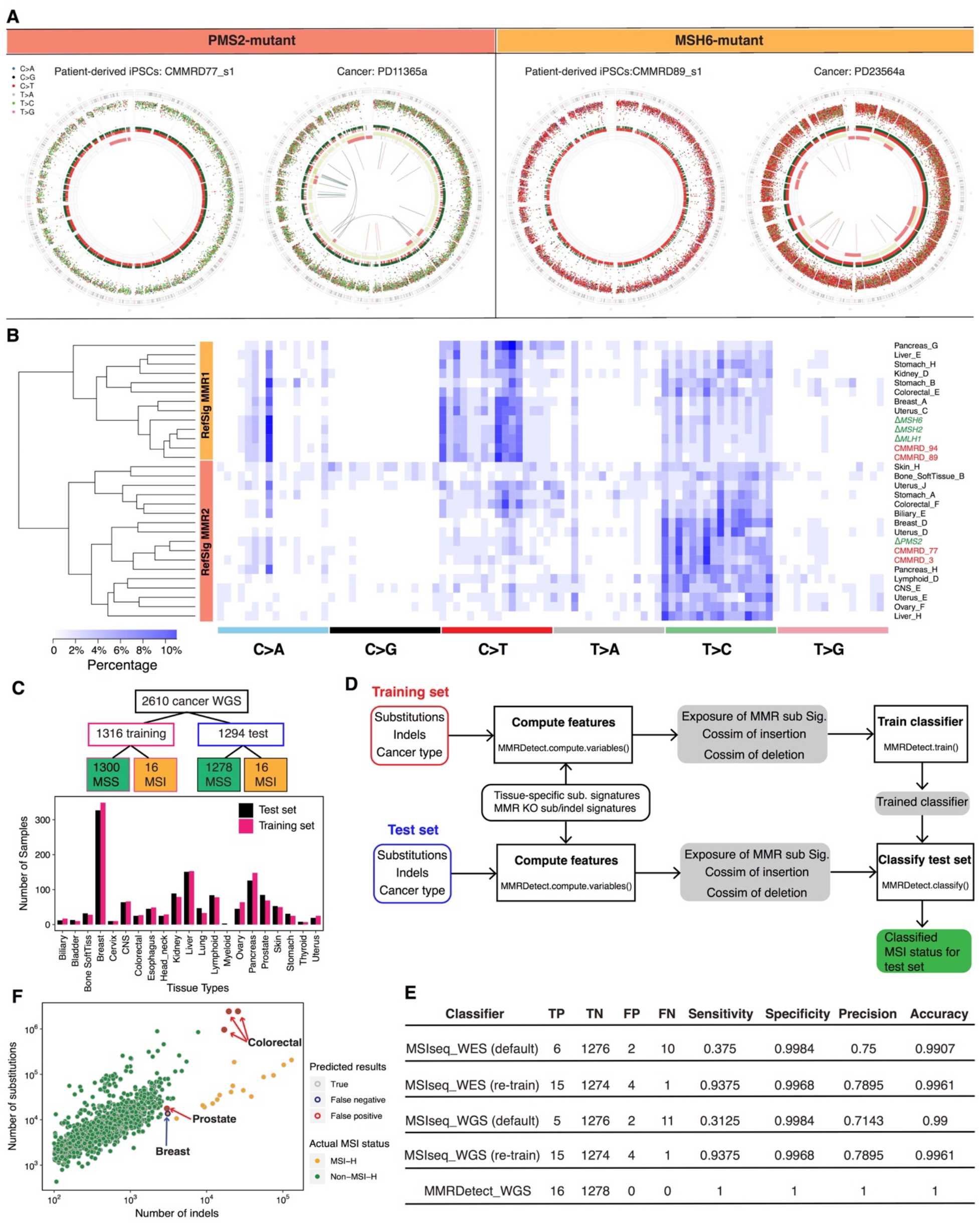
Gene-specific characteristics of mutational signatures of MMR-deficiency. (A) MMR knockouts demonstrate consistent gene-specificity regardless of model system, e.g., cancer *(in vivo)* and CMMRD patient-derived hiPSCs *(in vitro).* Whole-genome plots are shown for two patient-derived hiPSCs and two cancer samples. CMMRD77 is a *PMS2*-mutant patient. CMMRD89 is an *MSH6*-mutant patient. PD11365a and PD23564a are breast tumors with *PMS2* deficiency and *MSH2/MSH6* deficiency, respectively. Genome plots show somatic mutations including substitutions (outermost, dots represent six mutation types: C>A, blue; C>G, black; C>T, red; T>A, grey; T>C, green; T>G, pink), indels (the second outer circle, colour bars represent five types of indels: complex, grey; insertion, green; deletion other, red; repeat-mediated deletion, light red; microhomology-mediated deletion, dark red) and rearrangements (innermost, lines representing different types of rearrangements: tandem duplications, green; deletions, orange; inversions, blue; translocations, grey). See also Figure S14. (B) Hierarchical clustering of cancer-derived tissue-specific MMR signature and MMR knockout signatures. (C) Training and testing sample sets for MMR-deficiency classification. (D) Workflow of the MMRDetect. (E) Performance of MSIseq and MMRDetect. MSIseq was tested on WES and WGS data, using default classifier parameters and re-trained parameters. Data are shown in Tables S6 and S7. (F) Predicted results from MSISeq. Plot shows substitutions vs. indels for all samples tested.

Second, the predominance of C>A transversions could be explained by differential processing of 8-oxo-dGs (Figure 4C)^49,50^. The predominant C>A/G>T peak in MMR-deficient cells occurs at C<u>C</u>T>C<u>A</u>T/A<u>G</u>G>A<u>T</u>G followed by C<u>C</u>C>C<u>A</u>T/G<u>G</u>G>G<u>T</u>G and is distinct from the C>A/G>T peaks observed in Δ*OGG1* (Figure S11). However, we previously showed that there is a depletion of mutations at CC/GG sequence motifs for Δ*OGG1.* Intriguingly, the experimental data suggest that the 8-oxo-G:A mismatches can be repaired by MMR, preventing C>A/G>T mutations^51^. Furthermore, that G>T/C>A mutations of MMR-deficient cells occurred most frequently at the second G in 5’-TG_n_ (n>=3) in Δ*MLH1*, Δ*MSH2*, and Δ*MSH6* (Figures 3E and S12). This is consistent with previous reports^52^ of the classical imprint of guanine oxidation at polyG tracts where site reactivity in double-stranded 5’-TG^1^G^2^G^3^G^4^T sequence is reported as G^2^ > G^3^ > G^1^ > G^4^. These results implicate the activity of MMR in repairing 8-oxo-G:A mismatches at GG motifs that perhaps cannot be cleared by *OGG1* in BER. As for G>T leading strand bias, studies in yeast have demonstrated that an excess of 8-oxo-dG-associated mutations occurs during leading strand synthesis^53^. Furthermore, translesion synthesis polymerase η is also more error-prone when bypassing 8-oxo-dG on the leading strand^54^, which would result in increased 8-oxoG/A mispairs on the leading strand.

Third, we found that T>A transversions at A<u>T</u>T were strikingly persistent in MMR knockout signatures, although with modest peak size (<3% normalized signature, Figure 3A). Additional sequence context information revealed that T>A occurred most frequently at AA<u>T</u>TT or TT<u>T</u>AA, which were junctions of polyA and polyT tracts (Figure 3F)^55,56^. Moreover, the length of 5’- and 3’-flanking homopolymers influenced the likelihood of mutation occurrence: T>A transversions were one to two orders of magnitude more likely to occur when flanked by homopolymers of 5’polyA/3’polyT (A_n_T_m_) or 5’polyT/3’polyA (T_n_A_m_), than when there were no flanking homopolymeric tracts (Figure 3G).

Since polynucleotide repeat tracts predispose to indels due to replication slippage and are a known source of mutagenesis in MMR-deficient cells, we hypothesize that the T>A transversions observed at sites of abutting polyA and polyT tracts are the result of a ‘reverse template slippage’. In this scenario, the polymerase replicating across a mixed repeat sequence such as AAAAAATTTT in which the template slipped at one of the As would incorporate five instead of six Ts opposite the A repeat (red arrow pathway in Figure 3H). If at this point the template were to revert to its original correct alignment, this would give rise to an A/A mismatch that would result in a T>A transversion. If the slippage remained, this would give rise to a single nucleotide deletion, a characteristic feature of MMR-deficient cells known as microsatellite instability (MSI) (Figure 3B, indel signatures).

### Gene-specific characteristics of mutational signatures of MMR-deficiency

There are uncertainties regarding which of the cancer-derived signatures are truly MMR-deficiency signatures. It was suggested that SBS6, SBS14, SBS15, SBS20, SBS21, SBS26, and SBS44 were MMR-deficiency related^6^. In an independent analytical exercise, only two MMR-associated signatures were identified^24^, although variations of the signatures were seen in different tissue types^24^. An experimental process would help to obtain clarity in this regard^8–11^.

As described earlier, substitution patterns of Δ*MSH2*, Δ*MSH6*, and Δ*MLH1* showed enormous qualitative similarities to each other and were distinct from Δ*PMS2* (Figure 3A). We next expanded indel channels according to the length of polynucleotides, obtaining a higher resolution of MMR deficiency-associated indel signatures (Figure 3B). Δ*MSH2*, Δ*MSH6*, and Δ*MLH1* had very similar indel profiles, dominated by T deletions at increasing lengths of polyT tracts, with minor contributions of T insertions and C deletions. In contrast, Δ*PMS2* had similar proportions but different profiles between T insertions and deletions (Figures 3B and S8).

While the qualitative indel profiles of Δ*MSH2, ΔMSH6,* and Δ*MLH1* were very similar, their quantitative burdens were rather different (Figures 1E and S13). Δ*MLH1* and Δ*MSH2* had high indel burdens, while Δ*MSH6* had half the burden of indel mutagenesis. Substitution-to-indel ratios showed that Δ*MSH2, ΔPMS2,* and Δ*MLH1* produced similar amounts of substitutions and indels, while Δ*MSH6* generated nearly 2.5 times more substitutions than indels (Figure S13). This result is in-keeping with known protein interactions and functions: MSH2 and MSH6 form the heterodimer MutSα that addresses primarily base-base mismatches and small (1-2 nt) indels^46,57^. MSH2 can also heterodimerize with MSH3 to form the heterodimer MutS*β*, which does not recognize base-base mismatches, but can address indels of 1-15 nt^58^. This functional redundancy in the repair of small indels between MSH6 and MSH3 explains the smaller number of indels observed in Δ*MSH6* (Figure 1E) compared to Δ*MSH2* cells. This is consistent with the near-identical MSI phenotypes of Msh2^-/-^ and Msh3^-/-^; Msh6^-/-^ mice^59^.

Thus, there are clear qualitative differences between substitution and indel profiles of Δ*MSH2, ΔMSH6,* and Δ*MLH1* from Δ*PMS2.* To validate these two gene-specific experimentally-generated MMR knock-out signatures, we interrogated genomic profiles of normal cells derived from patients with inherited autosomal recessive defects in MMR genes resulting in Constitutional Mismatch Repair Deficiency (CMMRD), a severe, hereditary cancer predisposition syndrome characterized by an increased risk of early-onset (often pediatric) malignancies and cutaneous café-au-lait macules^60,61^. hiPSCs were generated from erythroblasts derived from blood samples of four CMMRD patients (two *PMS2* homozygotes and two *MSH6* homozygotes) and two healthy control^62^. hiPSC clones obtained were genotyped^62^. Expression arrays and cellomics-based immunohistochemistry were performed to ensure that pluripotent stem cells were generated (Methods). Parental clones were grown out to allow mutation accumulation, single-cell subclones were derived, and whole-genome sequenced (Figure S14A).

Gene-specificity of mutational signatures seen in CMMRD hiPSCs was virtually identical to those of the CRISPR-Cas9 knockouts and cancers (Figure 4A and S14B). The *PMS2* CMMRD patterns carried the same propensity for T>C mutations, the small contribution of C>T mutations and the single peak of C>A/G>T at C<u>C</u>T/A<u>G</u>G, as seen in Δ*PMS2,* and the *MSH6* CMMRD patterns carried the excess of C>T mutations with a very pronounced C>A/G>T at C<u>C</u>T/A<u>G</u>G similar to Δ*MLH1*, Δ*MSH2* and Δ*MSH6* clones (Figure S14C). Indel propensities seen in the knockout MMR clones were also reflected in the patient-derived cells (Figure S14D). Accordingly, gene-specificity of signatures generated in the experimental knockout system is well-recapitulated in an independent patient-derived cellular system of normal cells.

Furthermore, gene-specific MMR signatures were seen in the International Cancer Genome Consortium (ICGC) cohort of >2,500 primary WGS cancers^24^. Indeed, biallelic *MSH2/MSH6/MLH1* mutant tumors carried the same signature (RefSig MMR1) as Δ*MSH2* /Δ*MSH6*/Δ*MLH1* clones (Figure 4B). We also identified biallelic *PMS2* mutants in several cancers, including breast and ovarian cancers with mutation patterns (RefSig MMR2) that were indistinguishable from our experimentally-generated Δ*PMS2* signatures (Figure 4B).

### Informing classification of MMR-deficient tumors using experimental data

Algorithms to classify MMR-deficiency tumors have been developed using massively-parallel sequencing data^63–68^. These classifiers depend on detecting elevated tumor mutational burdens (TMB) or microsatellite instability (MSI). Mutation rates and cellular proliferative capacity are, however, variable between different tissues. New knowledge from our experimental data and awareness of tissue-specific signature variation (Figure 4B) led us to derive an MMR-deficiency classifier.

We used 2,610 published WGS primary cancers^69–71^ that had ≥200 substitutions and ≥100 indels (Methods) per tumor. Samples were randomly assigned into a training set (comprising 1,300 MMR-proficient microsatellite stable (MSS) and 16 MSI samples) or a test set (comprising 1,278 MSS and 16 MSI samples) (Figure 4C, Table S5). We trained a similar decision tree classifier to a widely-used MSI classifier, MSIseq^63^, using three new parameters based on the knowledge gained from these experiments, which we call MMRDetect: 1) the exposure of MMR-deficient substitution signatures; 2) the cosine similarity between repeat-mediated insertion profile of the tumor and that of MMR knockouts; and 3) the cosine similarity between repeat-mediated deletion profile of the tumor and that of MMR knockouts (Figure 4D, details of classifier development are provided in Methods). The performance of MMRDetect was compared with MSIseq^63^, using the same training and test data sets. Assessments were performed on whole-exome sequencing (WES) and WGS data using MSIseq’s default and re-trained classifiers (Figure 4E). MMRDetect achieved extremely high sensitivity and specificity (Figure 4E). MSIseq achieved relatively good performance following re-training of its classifier on this dataset, but this did not improve when using WGS data over WES data (Figure 4E), possibly because it does not increase the number of informative sites when scaled up to WGS. MSIseq has a higher likelihood of misclassifying non-MSI samples with high indel burden as MSI (false positives), and real MSI samples with low indel burden as non-MSI (false negatives) (Figure 4F). This is a generic problem for mutation burden-based MSI classifiers. For example, Nakagawa *et al.* used a cut-off of microsatellite mutation rate to define MSI samples^72^. According to their criterion, MSI samples with low indel burden would be missed because the cut-off is high, resulting in false-negative calls.

## Discussion

In standardized experiments performed in a diploid, non-transformed human stem cell model, biallelic gene knockouts that produce mutational signatures in the absence of administered DNA damage are indicative of genes that are important at maintaining the genome from intrinsic sources of DNA perturbations (Figure 5). We find signatures of substitutions and/or indels in nine genes: Δ*OGG1*, *DUNG*, Δ*EXO1*, Δ*RNF168*, Δ*MLH1*, Δ*MSH2*, Δ*MSH6*, Δ*PMS2*, and Δ*PMS1*, suggesting that proteins of these genes are critical guardians of the genome in non-transformed cells. Many gene knockouts did not show mutational signatures under these conditions. This does not mean that they are not important DNA repair proteins. There may be redundancy, or the gene may be crucial to the orchestration of DNA repair, even if itself is not imperative at directly preventing mutagenesis. For genes involved in double-strand-break (DSB) repair, hiPSCs may not be permissive for surviving DSBs to report signatures. Other genes may require alternative forms of endogenous DNA damage that manifest *in vivo* but not *in vitro,* for example, aldehydes, tissue-specific products of cellular metabolism, and pathophysiological processes such as replication stress. Likewise, for genes in the nucleotide excision repair pathway, bulky DNA adducts, whether exogenous (e.g., ultraviolet damage) or endogenous (e.g., cyclopurines and byproducts of lipid peroxidation) may be a pre-requisite before these compromised genes reveal associated signatures. While experimental modifications such as the addition of DNA damaging agents to increase mutation burden or using alternative cellular models, for example, cancer lines or cellular models of specific tissue-types, could amplify signal, they could also modify mutational outcomes, and that must be taken into consideration when interpreting data. Also, not all genes have been successfully knocked out in this endeavor and could have similarly important roles in directly preventing mutagenesis.

**Figure 5.**
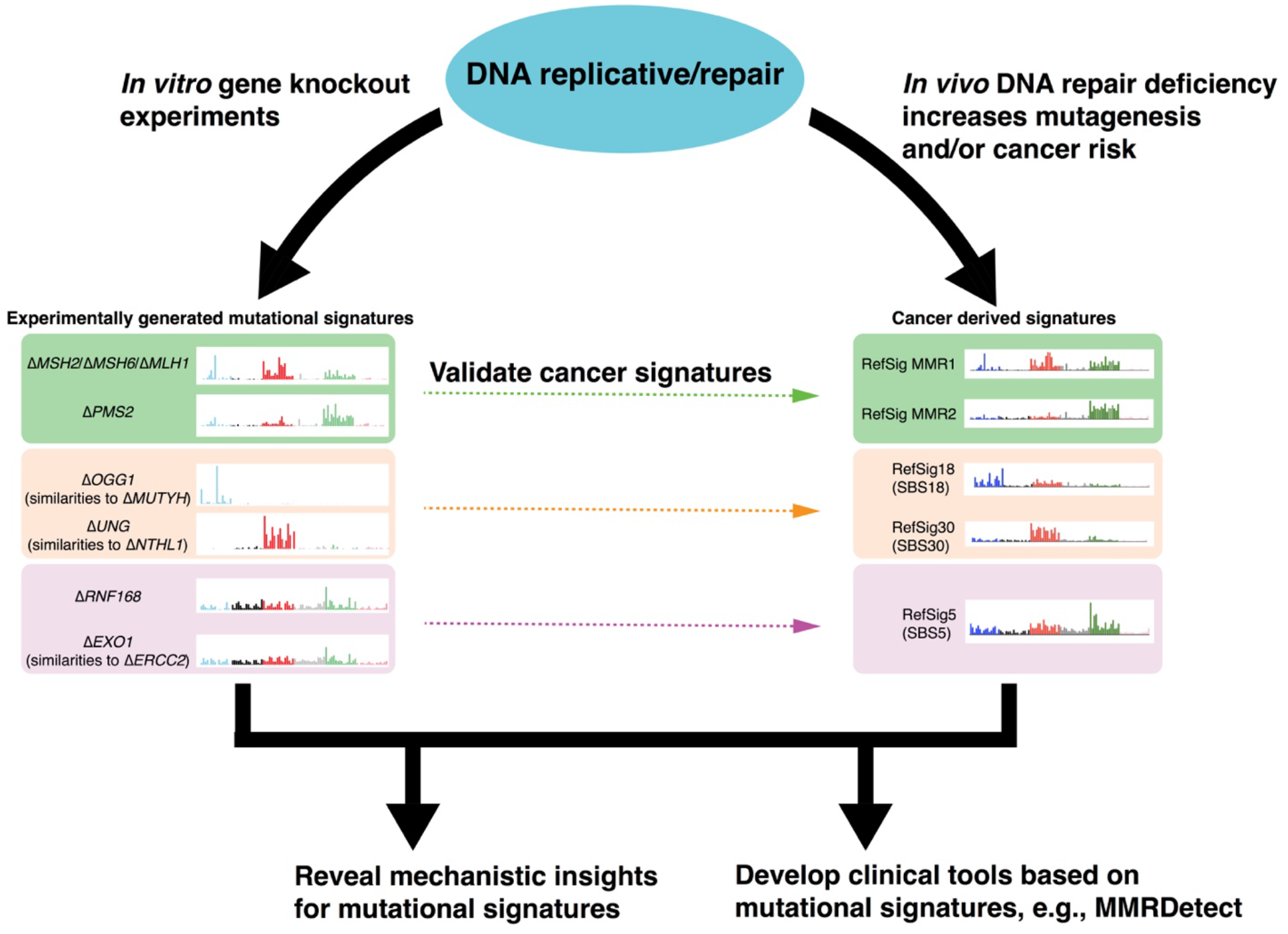
Impact of experimental validation of cancer-derived mutational signatures on biological understanding and development of clinical applications. Some genes (often involved in DNA repair pathways) which are important guardians against endogenous DNA damage under non-malignant circumstances, have been identified in this work. They help to validate and to understand the etiologies of cancer-derived mutational signatures. The biological insights help to drive the development of new genomic clinical tools to detect these abnormalities with greater accuracy and sensitivity across tumor types.

Upon detailed dissection of experimentally-generated signatures, we highlight interesting mutational insights, including how OGG1 and MMR proteins sanitize oxidized guanines at specific sequence motifs. By contrast, UNG maintains all cytosines from hydrolytic deamination reactions, irrespective of sequence context. Exhaustive assessment of DNA mismatches and their putative outcomes also revealed precise polymerase errors that are likely to be repaired by MMR, including misincorporation of T resulting in T>C transitions and misincorporation of G resulting G>A/C>T transitions by lagging strand polymerases. We also observe a T>A substitution at abutting poly-A and poly-T tracts that we postulate is due to a mechanism called reverse template slippage.

Of note, while it is known that 8-oxo-dGs can result in G>T mutations, our work demonstrates that the etiology of the culture-related signature and cancer-derived Signature 18, is mainly 8-oxo-dG. We highlight the importance of functional *EXO1* and *RNF168* in preventing Signature 5, a relatively ubiquitous signature characterized by T>C/A>G transitions. We define gene-specificities of signatures of MMR deficiency, prove that these are robust in normal, stem cells derived from patients with CMMRD, and also identify gene-specific signatures in human cancers.

Finally, unlike signatures of environmental mutagens that are historic, signatures of repair pathway defects are likely to be on-going in human cancer cells, and could serve as biomarkers of targetable abnormalities for precision medicine^13,14,18^ (Figure 5). This is important for pathways where there are selective therapeutic strategies available. These experiments led us to develop a more sensitive and specific mutational-signature-based assay to detect MMR deficiency, MMRDetect. Current TMB-based assays have reduced sensitivity to detect MMR deficiency because many tissues do not have high proliferative rates and may not meet the detection criteria of such assays. They may also falsely call MMR-deficient cases as MMR-proficient, because single components were used for measurement (e.g., indel burden or substitution count only). High mutational burdens can be due to different biological processes^73^. Consequently, assays based on burden alone are unlikely to be adequately specific. As a community, we are at the early stages of seeking experimental validation of mutational signatures. However, we hope that our approach, which leans on experimental data, provides a template for improving biological understanding of how mutational patterns arise, and that this, in turn, could help us propose improved tools for tumor characterization going forward.

## Acknowledgments

We thank the Wellcome Sanger Institute Cellular Genetics and Phenotyping facility for assistance, the CASM IT team, Julia Foreman and Grace Ping for assistance in carrying out and completing this project. We thank the COMSIG Consortium spearheaded by Professor Steve Jackson.

This work was funded by Cancer Research UK (CRUK) Advanced Clinician Scientist Award (C60100/A23916), Josef Steiner Cancer Research Award 2019, Medical Research Council (MRC) Grant-in-Aid to the MRC Cancer unit, CRUK Pioneer Award, BRC Cambridge core grant, Wellcome Strategic Award, and Wellcome Sanger Institute faculty funding. The work of T.I.R and J.S.C was funded by the CRUK Centre grant with reference number C309/A25144.

## Author contributions

Conceptualization: SNZ, BS; Writing: SNZ, XZ, GK, JJ, BS; Clinical sample collections: LS, GB, VPA, DR; Laboratory work: GK, KU, TIR, CAA, WB, CG; Data curation/Formal analysis: XZ, GK, ASN, AD, TIR, JSC, SNZ; Administration: RH, WB, JY.

## Materials & Correspondence

Further information and requests for materials may be directed to the corresponding author Serena Nik-Zainal.

## Methods

### Cell lines and culture

The human iPSC line used in this study is previously described (Kucab et al., 2019). The line was derived at the Wellcome Trust Sanger Institute (Hinxton, UK). The use of this cell line model was approved by Proportionate Review Sub-committee of the National Research Ethics (NRES) Committee North West – Liver-pool Central under the project “Exploring the biological processes underlying mutational signatures identified in induced pluripotent stem cell lines (iPSCs) that have been genetically modified or exposed to mutagens” (ref: 14.NW.0129). It is a long-standing iPSC line that is diploid and does not have any known driver mutations. It does carry a balanced translocation between chromosomes 6 and 8. It grows stably in culture and does not acquire a vast number of karyotypic abnormalities. This is confirmed through mutational and copy number assessment of the WGS data reviewed of all subclones.

Cell culture reagents were obtained from Stem Cell Technologies unless otherwise indicated. Cells were routinely maintained on Vitronectin XF-coated plates (10-15 ug/mL) in TeSR-E8 medium. The medium was changed daily, and cells were passaged every 4-8 days depending on the confluence of the plates using Gentle Cell Dissociation Reagent.

All cell lines were grown at 37°C, with 20% oxygen and 5% carbon dioxide in a humidified incubator, except for the pilot study in which the iPSCs knockouts were also grown under hypoxic condition (3% oxygen) as one of the experimental conditions (Supplementary Note 1). Cells were cultivated as monolayers in their respective growth medium and passaged every 3-4 days to maintain sub-confluence during the mutation accumulation step. All cell lines were tested negative for mycoplasma contamination using MycoAlertTM Mycoplasma Detection Kit and LookOut^®^ Mycoplasma PCR Detection Kit according to the manufacturers’ protocol.

### CMMRD patient sample collection

Four CMMRD patients were recruited at Doce de Octubre University Hospital, Spain, St George’s Hospital in London and Great Ormond Street Hospital under the auspices of the Insignia project. This included two PMS2-mutant patients and two MSH6-mutant patients. Table S8 shows the genotypes of these four patients. A healthy donor was recruited as control.

### Generation of DNA repair gene knockouts in human iPSCs

Biallelic DNA repair gene knockouts in human iPSCs were performed by the High Throughput Gene Editing team of Cellular Operations at the Sanger Institute, Hinxton, UK. These knockouts were generated based on the principles of CRISPR/Cas9-mediated HRD and NHEJ as described previously ^74^.

#### Generation of donor plasmids for precise gene targeting via HDR

All knockouts were generated using an established protocol that was found to minimize potential off-target effects ^74^. Briefly, the intermediate targeting vectors were generated for each gene using GIBSON assembly of the four fragments: pUC19 vector, 5’ homology arm, R1-pheS/zeo-R2 cassette and 3’ homology arm. Gene-specific homology arms were amplified by PCR from the iPSC gDNA and were either gel-purified or column-purified (QIAquick, QIAGEN). pUC19 vector and R1-pheS/zeo-R2 cassette were prepared as gel-purified blunt fragments (EcoRV digested). Fragments were assembled via GIBSON assembly reactions (Gibson Assembly Master Mix, NEB, E2611) according to the manufacturer’s instructions. Assembly reaction mix was transformed into NEB 5-alpha competent cells and clones resistant to carbenicillin (50 μg/mL) and zeocin (10 μg/mL) were analyzed by Sanger sequencing to select for correctly-assembled constructs. Sequence-verified intermediate targeting vectors were converted into donor plasmids via a Gateway exchange reaction. LR Clonase II Plus enzyme mix (Invitrogen, 12538120) was used to perform a two-way reaction exchanging only the R1-*pheSzeo*-R2 cassette with the pL1-EF1αPuro-L2 cassette as previously described^75^. The latter was generated by cloning synthetic DNA fragments of the EF1α promoter and puromycin resistance cassette into one of pL1/L2 vector ^75^. Following Gateway reaction and selection on yeast extract glucose (YEG) + carbenicillin agar (50 μg/mL) plates, correct donor plasmids were verified by capillary sequencing across all junctions.

#### Guide RNA design & cloning

For every gene knockout, two separate gRNAs targeting within the same critical exon of a gene were also selected. The gRNAs were selected using the WGE CRISPR tool ^76^ based on their off-target scores. Selected gRNAs were suitably positioned to ensure DNA cleavage within the exonic region, excluding any sequence within the homology arms of the targeting vector. To generate individual gene targeting plasmids, gene-specific forward and reverse oligos were annealed and cloned into BsaI site of either U6_BsaI_gRNA (kindly provided by Sebastian Gerety, unpublished). The gRNA sequences are listed in Table S9.

#### Delivery of KO-targeting plasmids, donor templates and Cas9, selection and genotyping

Human iPSCs were dissociated to single cells and nucleofected with Cas9-coding plasmid (hCas9, Addgene 41815), sgRNA plasmid and donor plasmid on Amaxa 4D-Nucleofactor program CA-137 (Lonza). Following nucleofection, cells were selected for up to 11 days with 0.25 μg/mL puromycin. Edited cells were expanded to ~70% confluency before subcloning. Approximately 1000 cells were subcloned onto 10 cm tissue culture dishes precoated with SyntheMAX substrate (Corning) at a concentration of 5 μg/cm^2^ to allow colony formation for 8-10 days until colonies are approximately 1-2 mm in diameter. Individual colonies were picked into U-bottom 96-well plates using a dissection microscope and a p20 pipette, grown to confluence and then replica plated. Once confluent, the replica plates were either frozen as single cells in 96-well vials or the wells were lysed for genotyping.

To genotype individual clones from a 96-well replica plate, cells were lysed and used for PCR amplification with LongAmp Taq DNA Polymerase (NEB, M0323). Insertion of the cassette into the correct locus was confirmed by visualizing on 1% E-gel (Invitrogen, G700801) PCR products generated by gene-specific (GF1 and GR1) and cassette specific primers (ER: TGATATCGTGGTATCGTTATGCGCCT and PF: CATGTCTGGATCCGGGGGTACCGCGTCGAG) for both 5’ and 3’ ends. We also confirmed single integration of the cassette by performing a qPCR copy number assay. To check the CRISPR site on the non-targeted allele, PCR products were generated from across the locus, using the same 5’ and the 3’ gene-specific genotyping primers. The PCR products were treated with exonuclease I and alkaline phosphatase (NEB, M0293; M0371) and Sanger sequenced to verify successful knockouts. Sequence reads and their traces were analysed and visualised on a laboratory information management system (LIMS)-2. For each targeted gene, two independently-derived clones with different specific mutations were isolated and studied further.

### Generation of iPSCs from Constitutional Mismatch Repair Deficiency (CMMRD) Patients

Peripheral blood mononuclear cells (PBMCs) isolation, erythroblast expansion, and IPSC derivation were done by the Cellular Generation and Phenotyping facility at the Wellcome Sanger Institute, Hinxton, according to Agu et al 2015^62^. Briefly, whole blood samples collected from consented CMMRD patients were diluted with PBS, and PBMCs were separated using standard Ficoll Paque density gradient centrifugation method. Following the PBMC separation, samples were cultured in media favoring expansion into erythroblasts for 9 days. Reprogramming of erythroblasts enriched fractions was done using non-integrating CytoTune-iPS Sendai Reprogramming kit (Invitrogen) based on the manufacturer’s recommendations. The kit contains three Sendai virus-based reprogramming vectors encoding the four Yamanaka factors, Oct3/4, Sox2, Klf4, and c-Myc. Successful reprogramming was confirmed via genotyping array and expression array.

### Proteomics analysis

Cell pellets were dissolved in 150 μL buffer containing 1% sodium deoxycholate (SDC), 100mM triethylammonium bicarbonate (TEAB), 10% isopropanol, 50mM NaCl and Halt protease and phosphatase inhibitor cocktail (100X) (Thermo, #78442) using pulsed probe sonication followed by boiling at 90 °C for 5 min. Aliquots containing 50 μg of total protein, measured with the Coomassie Plus Bradford Protein Assay (Pierce), were reduced with 5 mM tris-2-carboxyethyl phosphine (TCEP) for 1 h at 60 °C and alkylated with 10 mM Iodoacetamide (IAA) for 30 min in dark. Proteins were then digested with 75 ng/*μ*L trypsin (Pierce) overnight. The tryptic digests from the ATP2B4, EXO1, OGG1, PMS1, PMS2, RNF168 and UNG knock-out clones as well as three biological replicates of the parental cell line were labelled with the TMTpro 16plex reagents (Thermo) according to manufacturer’s instructions. The digests from MLH1, MSH2, MSH6 clones were subjected to label-free single-shot analysis. The TMTpro labelled peptides were fractionated with offline high-pH Reversed-Phase (RP) chromatography (XBridge C18, 2.1 x 150 mm, 3.5 μm, Waters) on a Dionex Ultimate 3000 HPLC system with 1% gradient. Mobile phase A was 0.1% ammonium hydroxide and mobile phase B was acetonitrile, 0.1% ammonium hydroxide. LC-MS analysis was performed on the Dionex Ultimate 3000 system coupled with the Orbitrap Lumos Mass Spectrometer (Thermo Scientific). Selected TMTpro peptide fractions were loaded to the Acclaim PepMap 100, 100 μm × 2 cm C18, 5 μm, 100 A trapping column and were analyzed with the EASY-Spray C18 capillary column (75 μm × 50 cm, 2 μm). Mobile phase A was 0.1% formic acid and mobile phase B was 80% acetonitrile, 0.1% formic acid. The TMTpro peptide fractions were analyzed with a 90 min gradient from 5%-38% B. MS spectral were acquired with mass resolution of 120 k and precursors were isolated for CID fragmentation with collision energy 35%. MS3 quantification was obtained with HCD fragmentation of the top 5 most abundant CID fragments isolated with Synchronous Precursor Selection (SPS) and collision energy 55% at 50k resolution. For the label-free experiments, peptides were analyzed with a 240 min gradient and HCD fragmentation with collision energy 35% and ion trap detection. Database search was performed in Proteome Discoverer 2.4 (Thermo Scientific) using the SequestHT search engine with precursor mass tolerance 20 ppm and fragment ion mass tolerance 0.5 Da. TMTpro at N-terminus/K (for the labelled samples only) and Carbamidomethyl at C were defined as static modifications. Dynamic modifications included oxidation of M and Deamidation of N/Q. The Percolator node was used for peptide confidence estimation and peptides were filtered for q-value < 0.01. All spectra were searched against reviewed UniProt human protein entries. Only unique peptides were used for quantification. The results of proteomics analysis are provided in Table S10.

### Proliferation assay

Cells were seeded at 5,500 per well on 96-w plates. Measurements were taken at 24 h intervals post-seeding over a period of 5 days according to manufacturer’s instructions. Briefly, plates were removed from the incubator and allowed to equilibrate at room temperature for 30 minutes, and equal volume of CellTiter-Glo reagent (Promega) was added directly to the wells. Plates were incubated at room temperature for 2 minutes on a shaker and left to equilibrate for 10 minutes at 22°C before luminescence was measured on PHERAstar *FS* microplate reader. Luminescence readings were normalized and presented as relative luminescence units (RLU) to time point 0 (t_0_). Figure S15 shows the statistics of 6 replicates for each time point per indicated knockout lines. Error bars show standard error of the mean. Doubling time was calculated based on replicate-averaged readings on the linear portion of the proliferation curve (exponential phase) using formula:

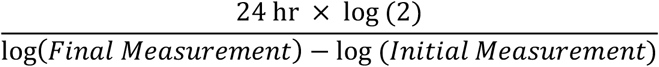

### Genomic DNA extraction and WGS

Samples were quantified with Biotium Accuclear Ultra high sensitivity dsDNA Quantitative kit using Mosquito LV liquid platform, Bravo WS and BMG FLUOstar Omega plate reader and cherrypicked to 500ng/120μl using Tecan liquid handling platform. Cherrypicked plates were sheared to 450bp using a Covaris LE220 instrument. Post-sheared samples were purified using Agencourt AMPure XP SPRI beads on Agilent Bravo WS. Libraries were constructed (ER, A-tailing and ligation) using ‘Agilent Sureselect kit’ on an Agilent Bravo WS automation system. KapaHiFi Hot start mix and IDT 96 iPCR tag barcodes were used for PCR set-up on Agilent Bravo WS automation system. PCR cycles include 6 standard cycles: 1) Incubate 95C 5 mins; 2) Incubate 98C 30 secs; 3) Incubate 65C 30 secs; 4) Incubate 72C 1 min; 5) Cycle from 2, 5 more times; 6) Incubate 72C 10 mins. Post PCR plate was purified using Agencourt AMPure XP SPRI beads on Beckman BioMek NX96 liquid handling platform. Libraries were quantified with Biotium Accuclear Ultra high sensitivity dsDNA Quantitative kit using Mosquito LV liquid handling platform, Bravo WS and BMG FLUOstar Omega plate reader, then pooled in equimolar amounts on a Beckman BioMek NX-8 liquid handling platform and finally normalized to 2.8 nM ready for cluster generation on a c-BOT and loading on requested Illumina sequencing platform. Pooled samples were loaded on the X10 using 150 PE run length, sequenced to ~25X coverage. The details of sequence coverage for all clones and subclones are provided in Table S2.

### Alignment and somatic variant-calling

Short reads were aligned to human reference genome GRCh37/hg19 assembly using the BWA-MEM algorithm^77^. Three algorithms, CaVEMan (http://cancerit.github.io/CaVEMan/)^78^, Pindel (http://cancerit.github.io/cgpPindel)^79^ and BRASS (https://github.com/cancerit/BRASS) were used to call somatic substitutions, indels and rearrangements in all subclones, respectively.

### Assurance of knockout state using WGS data

First, we examined whether there were CRISPR-Cas9 off-target effects by seeking relevant mutations in other DNA repair genes besides the genes of interest. We also searched for potential off-target sites based on gRNA target sequences using COSMID^80^ and confirmed that there were no off-target hits in knockouts that generated mutational signatures (Table S11). We confirmed chromosome copy number in all subclones remained stable and unchanged from their parent. Second, we confirmed that there are frameshift indels near the gRNA targeted sequence in the genes of interest for all knockout subclones. One *UNG* knockout was found to be heterozygous and was excluded in the downstream analysis. Third, we checked mislabelled samples by examining the shared mutations between subclones. Subclones originally derived from the same parental knockout clone would share some mutations, in contrast to subclones from different knockouts. Consequently, one Δ*PRKDC,* one Δ*TP53* and two Δ*NBN* subclones were removed from downstream analysis. Fourth, variant allele fraction (VAF) distribution for each knockout subclone was examined. VAF>=0.4 was used as a cut-off for determination of whether the subclone was derived from a single-cell. When contrasting mutation burden between subclones, we only selected subclones that were derived from single-cells, cultured for 15 days. Shared mutations among subclones were removed to obtain *de novo* somatic mutations accumulated after knocking out the gene of interest. Table S2 summarizes the number of *de novo* mutations (substitutions and indels) for all subclones.

### Determination of gene knockout-associated mutational signatures

An intrinsic background mutagenesis exists in normal cells grown in culture. Knocking out a DNA repair gene that is involved in repairing endogenous DNA damage may result in increased unrepaired DNA damage and, thereby result in mutation accumulation with subsequent rounds of replication. Whole-genome sequencing of these knockouts can detect the mutations that occur as a result of being a specified knockout. If the mutation burden and the mutational profile of a knockout is significantly different from the control subclones which have only the background mutagenesis, it is most likely that there is gene knockout-associated mutagenesis. Based on this principle, our approach to identify gene knockout-associated mutational signature involved three steps: 1) we determined the background mutational signature; 2) we determined the difference between the mutational profile of knockout and background mutation profiles. 3) we removed the background mutation profile from mutation profile of the knockout subclone.

Substitution profiles were described according to the classical convention of 96 channels: the product of 6 types of substitution multiplied by 4 types of 5’ base (A,C,G,T) and 4 types of 3’ base (A,C,G,T). Indel profiles were described by type (insertion, deletion, complex), size (1-bp or longer) and flanking sequence (repeat-mediated, microhomology-mediated or other) of the indel. Here, we used two sets of indel channels. Set one contains 15 channels: 1bp C/T insertion at short repetitive sequence (<5 bp), 1bp C/T insertion at long repetitive sequence (>=5 bp), long insertions (> 1bp) at repetitive sequences, microhomology-mediated insertions, 1bp C/T deletions at short repetitive sequence (<5 bp), 1bp C/T deletions at long repetitive sequence (>=5 bp), long deletions (> 1bp) at repetitive sequences, microhomology-mediated deletions, other deletion and complex indels (Figure S9). Set two contains 45 channels, in which the 1 bp C/T indels at repetitive sequences are further expanded according to the exact length of the repetitive sequences (Figure 3B). Indel channel set one was applied to all knockout subclones, whilst channel set two was only applied to four MMR gene knockouts (Δ*MLH1*, Δ*PMS2*, Δ*MSH2*, Δ*MSH6*) to obtain a higher resolution of mutational signatures of MMR gene knockouts.

#### Identifying background signatures

The mutational profile of control subclones were used to determine background mutagenesis. Aggregated substitution profiles of all control subclones (Δ*ATP2B4*) were used as the background substitution mutational signature. Aggregated indel profiles of all subclones containing = 8 indels were used as the background indel mutational signature.

#### Distinguishing mutational profiles of control and gene-edited subclone profiles

Signal-to-noise ratio affects mutational signature detection. In this study, ‘noise’ is largely background mutagenesis. The averaged mutation burden caused by the background mutagenesis in control cells for substitution and indels are around 150 and 10, with standard deviation of 10 and 1.4, respectively. ‘Signal’ represents the elevated mutation burden caused by gene knockouts. The averaged mutation burden in knockouts range from 63 to 2360 for substitution, and 0 to 2122 for indels after 15 days in culture, as shown in Table S2.

The costs associated with whole genome sequencing is prohibitive, thus we have 2-4 subclones per knockout. The intrinsic fluctuation of detected mutation burden in each sample and the limited subclone numbers impose a greater uncertainty in mutational signature detection. Thus, to distinguish high-confidence mutational signatures from noise, we employed three different methods.

First, we evaluated the similarity of mutational profile between control and each gene knockout. According to the mutational profile of control subclones, 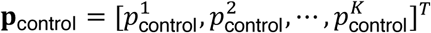, for a given number of mutations *N* (0 < *N* < 10000), one could generate *L* bootstrapped samples:

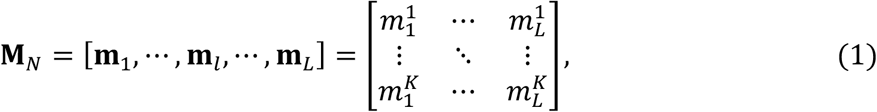

where 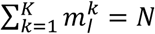. One can calculate the cosine similarities (*s_l_*) between bootstrapped control samples (**m**_*l*_) and experimentally-obtained control profile (**p**_control_) to obtain a distribution of cosine similarities *P*(*S*):

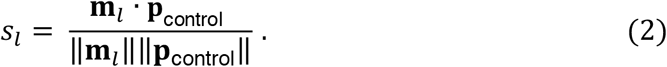

We can then calculate the cosine similarity (*S*_knockout_) between control profile (**p**_control_) and knockout profile (**p**_knockout_). As shown in Figures 1C and 1D, when the mutation count is low, the bootstrapped samples are less similar to the actual control profile than the bootstrapped samples with higher mutation count. Comparing *S*_knockout_ and *P*(*S*) at a given mutation number, *N*_knockout_, one could identify which gene knockouts having distinct mutational profiles from the control (p value of *S*_knockout_ is less than 0.01 in *P*(*S*)).

Second, we used contrastive principal component analysis (cPCA) ^21^, which efficiently identified directions that were enriched in the knockouts relative to the background through eliminating confounding variations present in both (Figure S4A), to recognize gene knockout-specific patterns from background signature.

Third, we used t-Distributed stochastic neighbor embedding (t-SNE) ^22^, which is a visualization technique for viewing pairwise similarity data resulting from nonlinear dimensionality reduction based on probability distributions. In t-SNE implementation, mutational profiles that are similar to each other were plotted nearby each other, whereas profiles that are dissimilar are plotted distantly in a 2D space (Figure S4B).

#### Subtraction of the background mutational signature from knockout mutation profile

The experiment-associated mutational signature can then be obtained by subtracting the background mutational signature from the mutational profile of treated subclones through quantile analysis. First, one can generate a set of bootstrap samples of each treated subclone in order to determine the distribution of mutation number for each channel. According to the distribution, the upper and lower boundaries (e.g., 99% CI) for each channel can be identified. Then, based on the background mutational signature and averaged mutation burden (as initial value), one can construct bootstrapped background profiles, and subtract it from the centroid of bootstrap subclone samples. Due to data noise, some channels may have negative values, in which case, the negative values are set to zero. Occasionally, the number of mutations in a few channels will fall outside the lower boundary after removing the background profile. To avoid negative values, the background mutation pattern is maintained but burden is scaled down through an automated iterative process.

### Topography analysis of signatures

#### Strand bias

Reference information of replicative strands and replication-timing regions were obtained from Repli-seq data of the ENCODE project (https://www.encodeproject.org/) ^81^. The transcriptional strand coordinates were inferred from the known footprints and transcriptional direction of protein coding genes. First, for a given mutational signature, one could calculate the ‘expected’ ratio of mutations between transcribed and non-transcribed strand, or between lagging and leading strands, according to the distribution of trinucleotide sequence context in these regions. Second, the ‘observed’ ratio of mutations between different strands can be identified through mapping mutations to the genomic coordinates of all gene footprints (for transcription) or leading/lagging regions (for replication). Third, all mutations were orientated towards pyrimidines as the mutated base (as this has become the convention in the field). This helped denote which strand the mutation was on. Fourth, the level of asymmetry between different strands was measured by calculating the odds ratio of mutations occurring on one strand (e.g., transcribed or leading strand) vs. on the other strand (e.g., non-transcribed or lagging strand).

### MMRDetect algorithm

We trained an MSI classifier based on mutational signatures (MMRDetect) using the same decision tree framework that previously utilized by MSIseq^63^. We obtained published wholegenome somatic mutation data and MSI statuses from 2610 tumors from three different studies^69–71^. These tumors involve 21 cancer types. All 2610 cancers have ≥200 substitutions and ≥100 indels and also have both substitution and indels in exons. 2610 cancers were randomly split into a training data set and a test data set by using the R function sample(). The training data set had 1300 MSS and 16 MSI samples. The test data set had 1278 MSS and 16 MSI samples (Table S5). To ensure there was no tissue-specific bias in sample assignment, we confirmed that each cancer type had similar sample numbers in training and test data sets. We fitted tissue-specific substitution signatures to each tumor using an R package (signature.tools.lib). The sum of exposures of MMR1 and MMR2 was used as a feature in MMRDetect. The cosine similarities between the profiles of repeat-mediated insertions/deletions of cancer samples and those of MMR gene knockouts were calculated and used as the other two features in MMRDetect. Tables S6 and S7 show calculated parameters of 2610 tumors for MSIseq and MMRDetect, respectively. The decision tree algorithm (function J48()) provided in R package RWeka^82^ was employed as the framework of MMRDetect.

We used several metrics to evaluate different classifiers. First, we counted the numbers of true positive (TP), true negative (TN), false positive (FP) and false negative (FN) samples generated by each classifier. Then we calculated sensitivity, specificity, precision and accuracy for each classifier using the following equations:

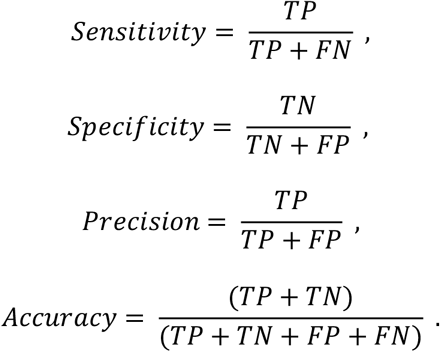

### Rational of choosing parameters for the classifier

We fitted tissue-specific substitution signatures to each sample, thereby acquiring exposures to MMR deficiency signatures, *E*_MMR_. MSI samples had increased exposures of MMR1 and MMR2 signatures (p-value = 2.7×10^-17^, Mann-Whitney test, function wilcox.test() in R) (Figure S16). However, a few non-MSI samples were also assigned with MMR signatures, possibly due to overfitting. Thus, to improve specificity, we examined indel profiles of all samples. We calculated cosine similarities between repeat-mediated indels profiles of cancer samples and MMR knockout indel signatures for insertion (*S*_ins_)and deletion (*S*_del_), respectively. MSI samples and MMR gene knockouts showed near-identical profiles of repeat-mediated deletions and insertions, whilst MSS samples had lower cosine similarities for deletions (Figure S16, p-value = 9.3×10^-12^, Mann-Whitney test) and insertions (Figure S16, p-value = 5.5×10^-8^, Mann-Whitney test). Indeed, a distinct separation pattern can be observed by the combination of these three features.

#### Other software used in the study

IntersectBed ^83^ was used to identify mutations overlapping certain genomic features. All statistical analysis were performed in R ^84^. All plots were generated by ggplot2 ^85^.

#### Data and software availability

Raw sequence files are to be deposited at the European Genome-phenome Archive with accession numbers EGAS00001000800 and EGAS00001000874. Mutation calls have been deposited at Mendeley and the link will be provided once the manuscript is accepted. hiPSCs can be obtained directly from the authors. Code for analysis will be available on GitHub once the manuscript is accepted.

The curated data will be available for general browsing from our reference Mutational Signature website, SIGNAL (https://signal.mutationalsignatures.com).

## Extended data

Supplementary Information. Figures S1–S16. Table S8

Table S1. List of DNA repair genes targeted.

Table S2. List of 173 gene knockout subclones.

Table S3. Substitution catalogue of subclones.

Table S4. Mutation rates of knockout-generated mutational signatures.

Table S5. List of 2610 cancer samples.

Table S6. Calculated input parameters for MSIseq.

Table S7. Calculated input parameters for MMRDetect.

Table S9. gRNA sequences for DNA repair gene knockouts in human iPSCs.

Table S10. Results of proteomics analysis.

Table S11. COSMID output: a ranked-list of potential off-target sites of gRNA sequences.

## Supplementary Figures

**Figure S1.**
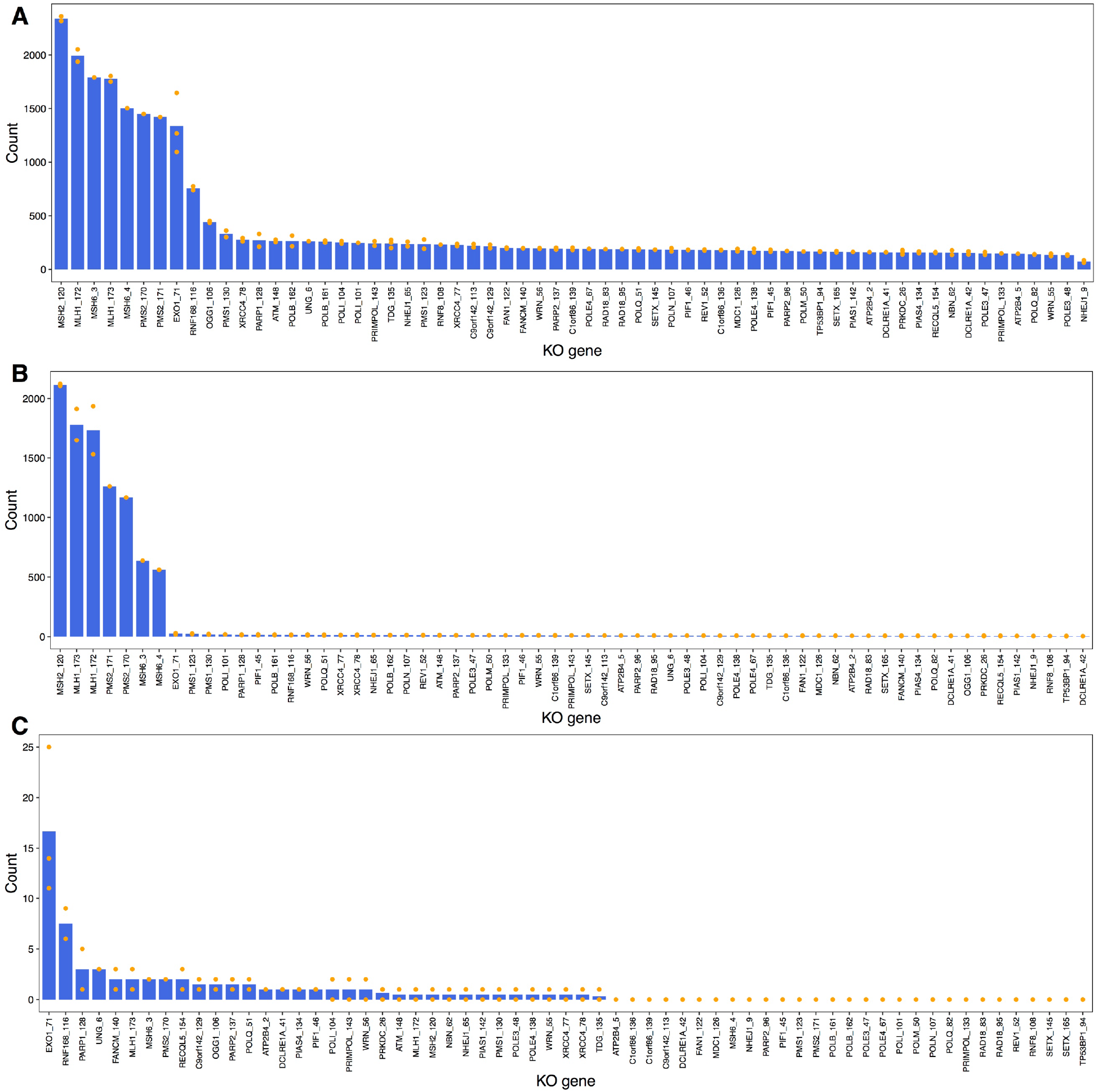
Substitution (A), indel (B) and double substitution (C) burden of gene knockout subclones. Only clonal daughter subclones cultured for 15 days were included in all comparative analyses.

**Figure S2.**
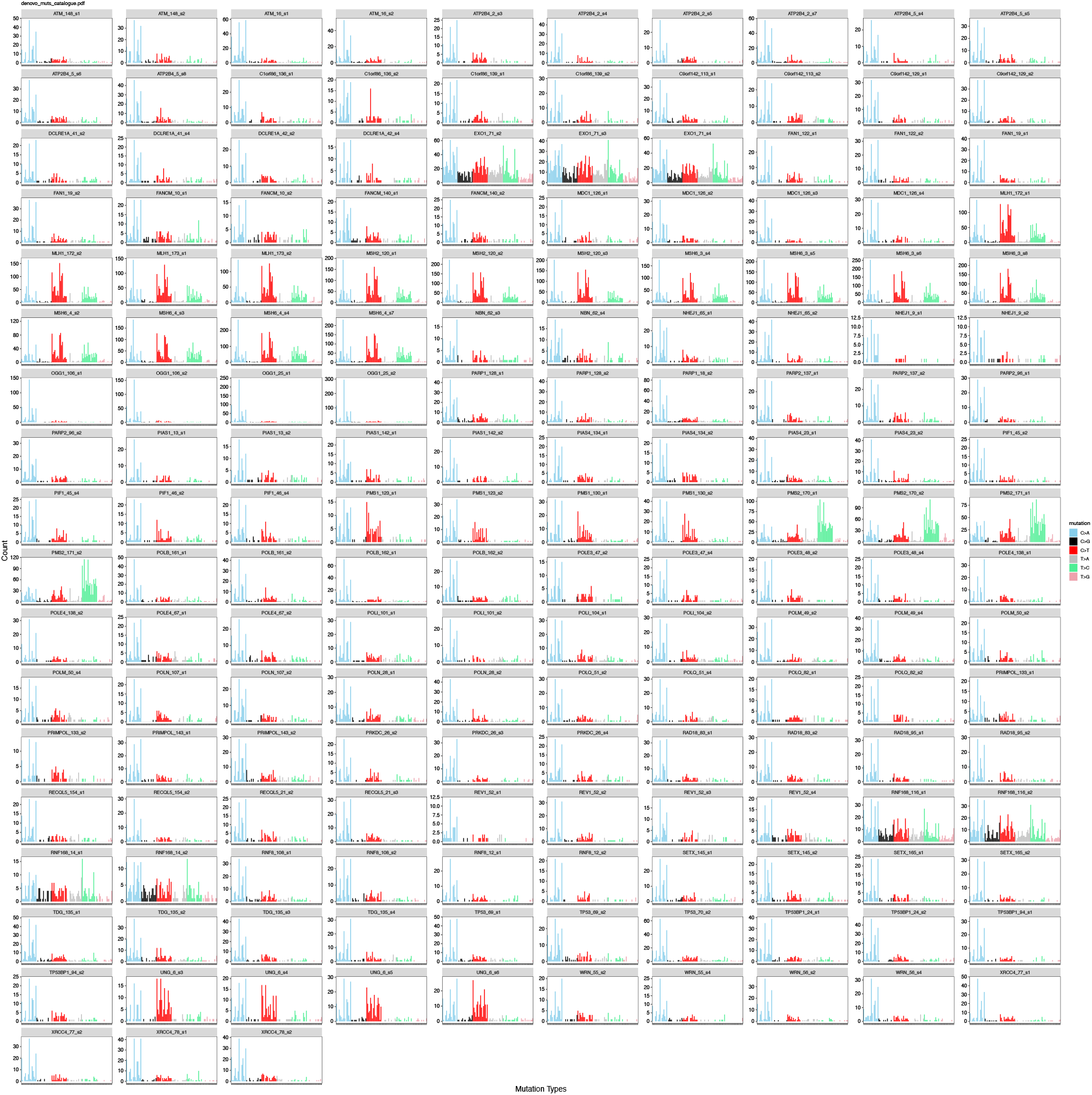
96-channel substitution mutation profiles of 173 gene knockout subclones.

**Figure S3.**
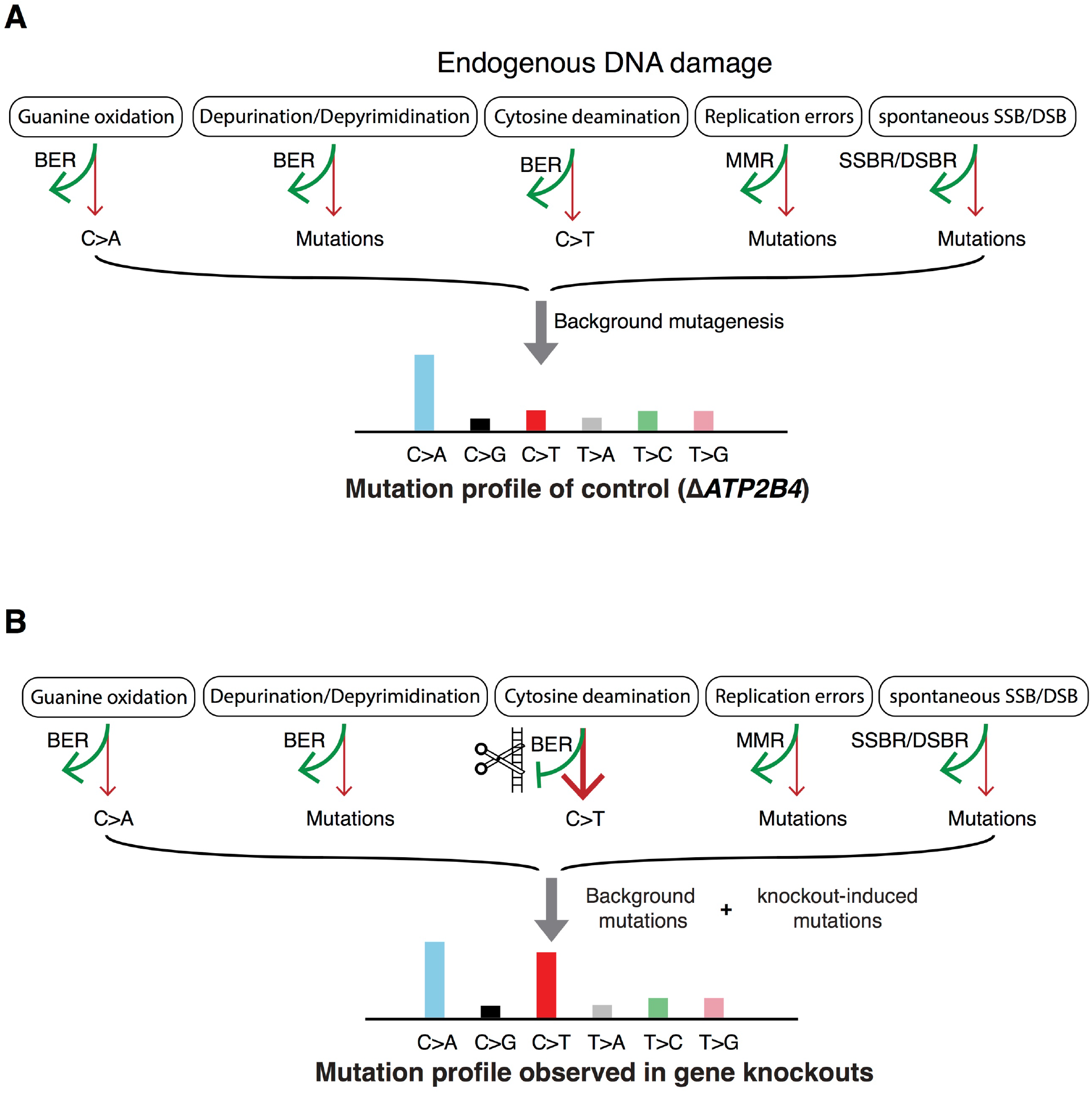
Schematic depicting the principle of detecting mutational consequences of knockouts in the absence of added external DNA damage. (A) Potential components of background signature. (B) Possible mutational consequences of the DNA repair gene knockouts for proteins that are critical mitigators of mutagenesis.

**Figure S4.**
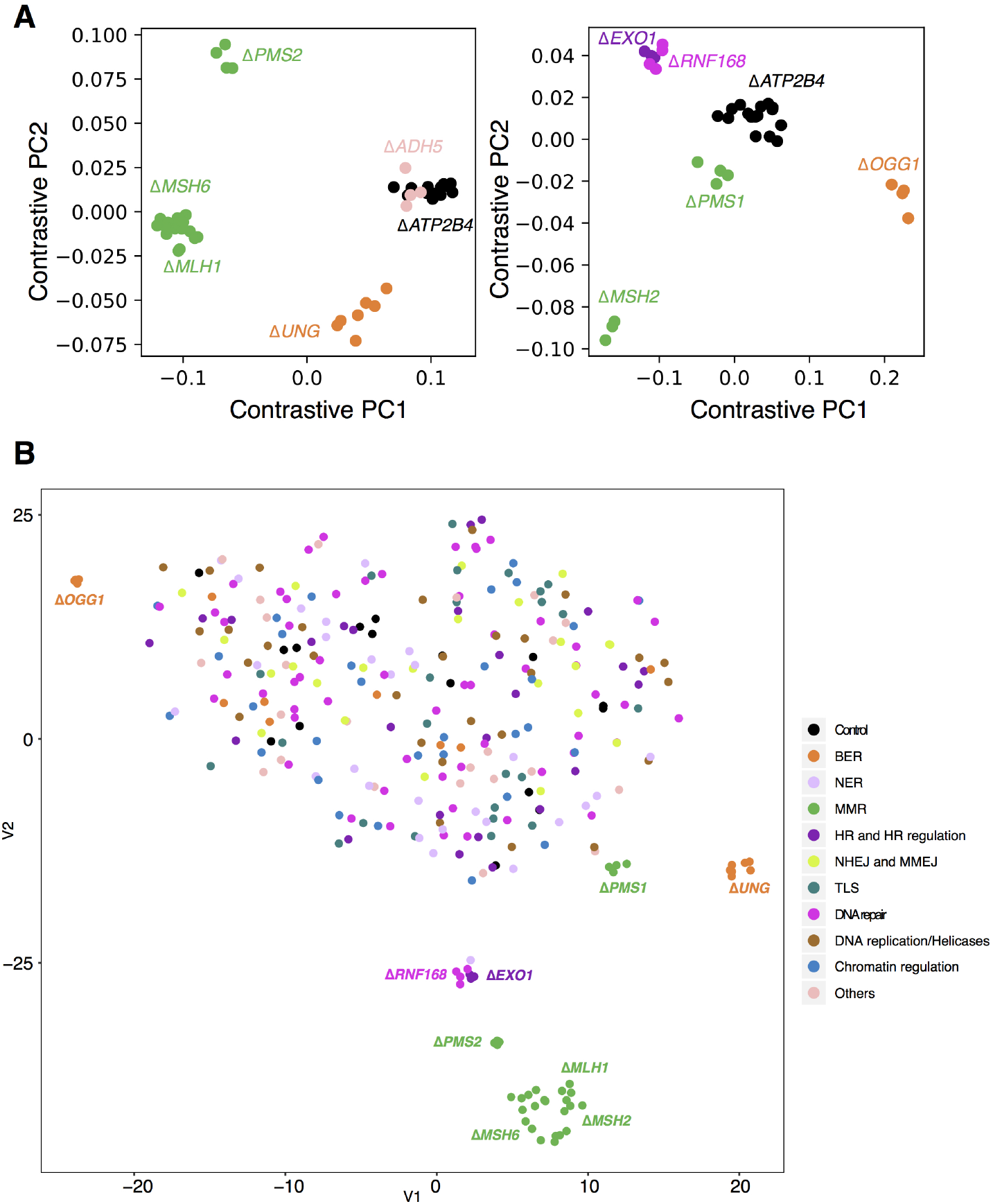
Results of contrastive principal component analysis and t-SNE. (A) Contrastive principal component analysis (cPCA) was employed to discriminate knockout profiles from control profiles (Δ*ATP2B4*). Each figure contains six different genes. Nine gene knockouts separate from the controls. Using this method, Δ*ADH5* did not separate clearly from Δ*ATP2B4,* indicative of either having no signature or a weak signature. Dot colour indicate the repair/replicative pathway that each gene is involved in: black - control; green - MMR; orange – BER; dark purple – HR and HR regulation; light purple - checkpoint. (B) The t-SNE algorithm was applied to discriminate the mutational profiles of gene knockouts from those of control knockouts. Gene knockouts that produce mutational signatures separate clearly from control subclones and other knockouts which do not have signatures. Subclones of the gene knockouts which produce signatures are clustered together, indicating consistency between subclones.

**Figure S5.**
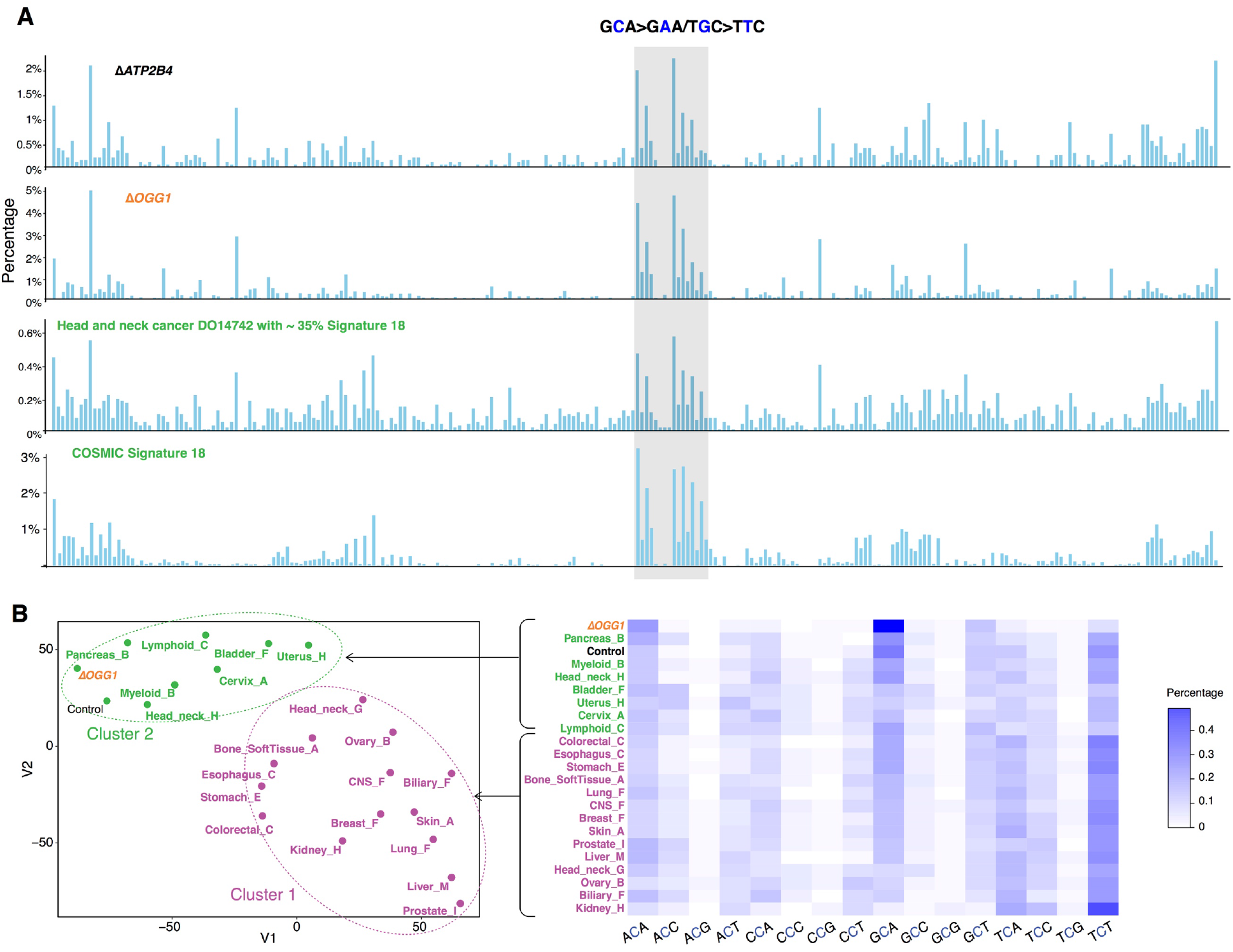
(A) Relative mutation frequency of G>T/C>A in 256 possible channels which take two adjacent bases 5’ and 3’ of each mutated base (4×4×4×4=256) for Δ*ATP2B4*, Δ*OGG1*, a head and neck cancer with strong Signature 18 and COSMIC Signature 18. (B) Left: tSNE plot of tissuespecific mutational signature 18. Two groups are featured with predominant peaks at T<u>G</u>C>T<u>T</u>C/G<u>C</u>A>G<u>A</u>A (highlighted in green) and A*G*A>A<u>T</u>A/T<u>C</u>T>T<u>A</u>T (highlighted in purple), respectively. Right: heatmap of 21 tissue-specific mutational signatures at C>A. We compared experimental signatures to previously published cancer-derived signatures, focusing on 21 tissuespecific variations of Signature 18^18^. Interestingly, we found two distinct groups of Signature 18. Signatures of Δ*OGG1*, cellular models and signatures derived from head and neck tumors, pancreas, myeloid, bladder, uterus, cervix, lymphoid tumors were most similar to each other, with the predominant G>T/C>A peak at T<u>G</u>C>T<u>T</u>C/G<u>C</u>A>G<u>A</u>A. By contrast, an alternative version of this signature with a predominant G>T/C>A peak at A<u>G</u>A>A<u>T</u>A/T<u>C</u>T>T<u>A</u>T was noted in colorectal, esophagus, stomach, bone, lung, CNS, breast, skin, prostate, liver, head and neck tumors (Signature Head_neck_G), ovary, biliary and kidney cancers. Indeed, there are many types of oxidative species which could fluctuate between tissues, variably affecting trinucleotides resulting in the variation observed in Signature 18.

**Figure S6.**
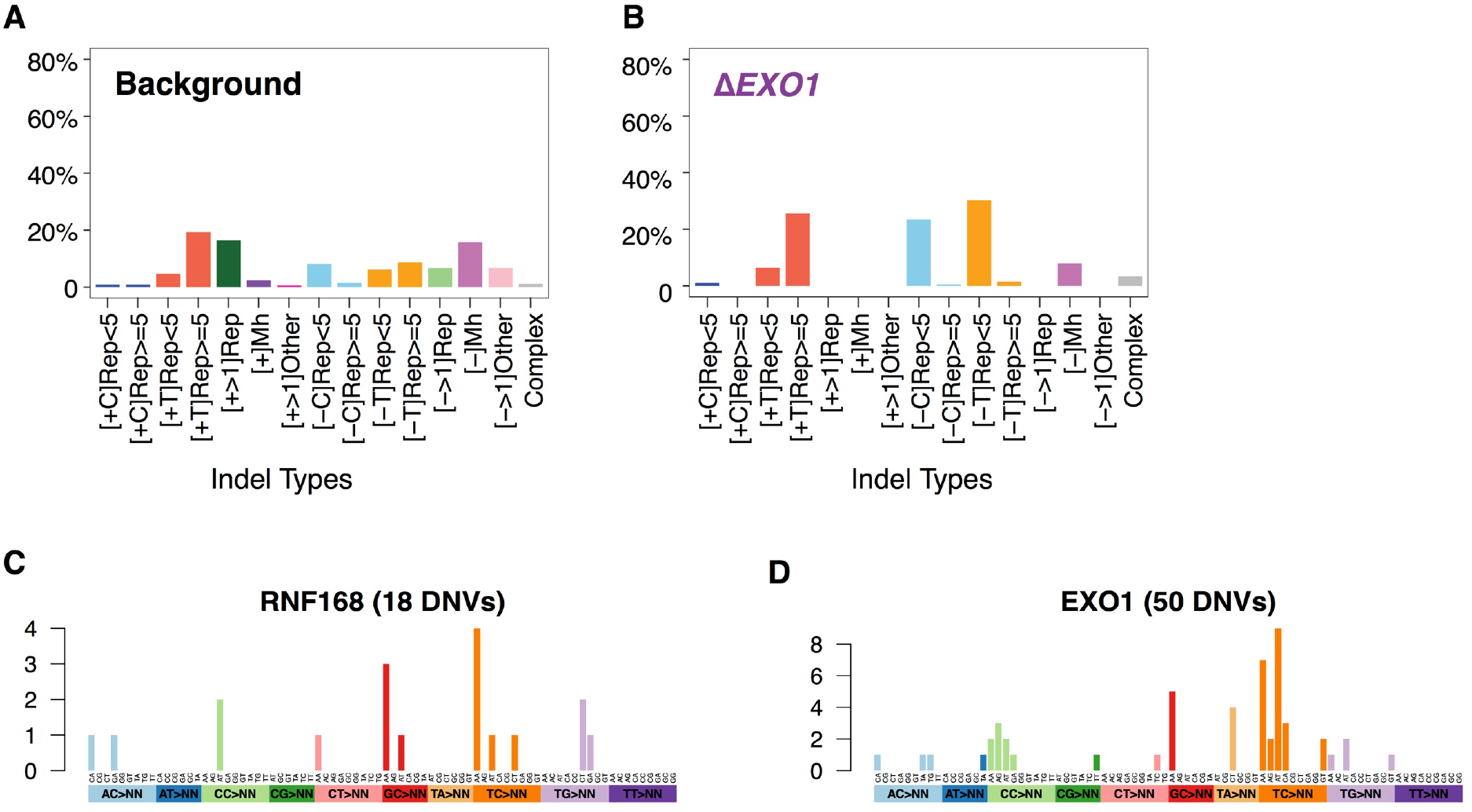
(A) Background indel signature. (B) Indel signature of Δ*EXO1.* (C) Aggregated double substitution profile of Δ*RNF168*. (D) Aggregated double substitution profile of Δ*EXO1*.

**Figure S7.**
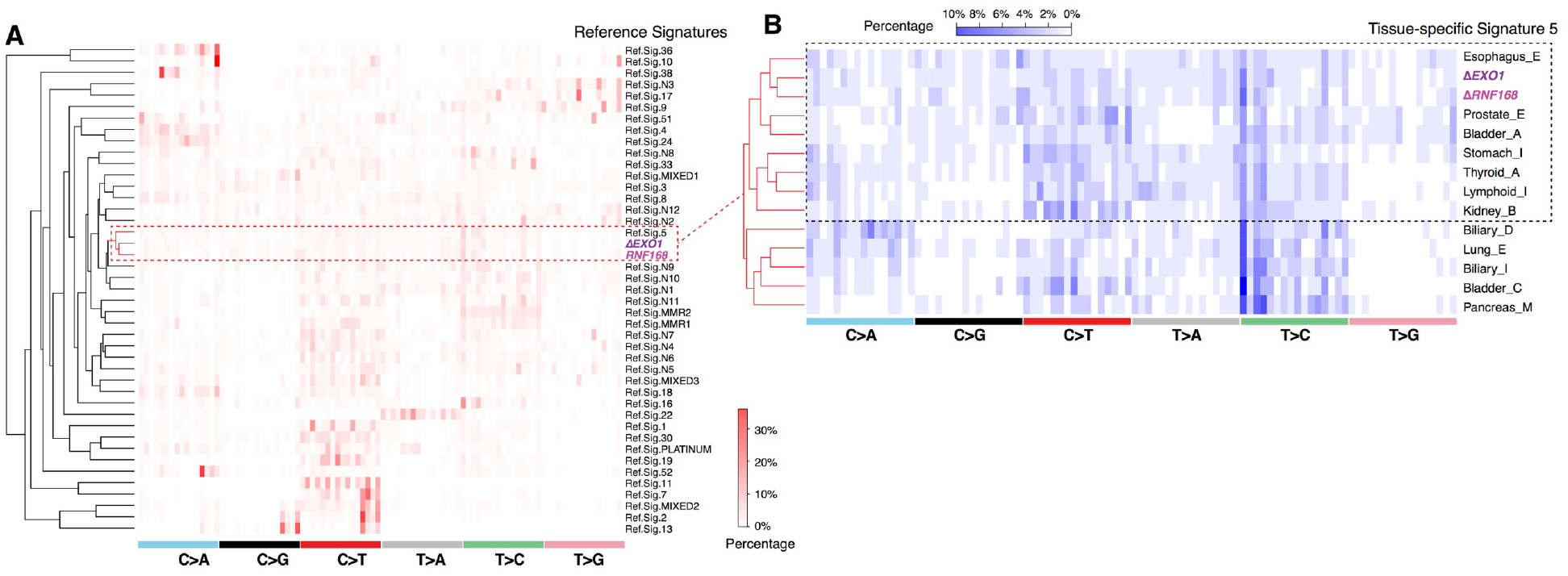
(A) Hierarchical clustering of cancer-derived reference signatures with Δ*EXO1* and Δ*RNF168* signatures. (B) Hierarchical clustering of tissue-specific signature 5 with Δ*EXO1* and Δ*RNF168* signatures.

**Figure S8.**
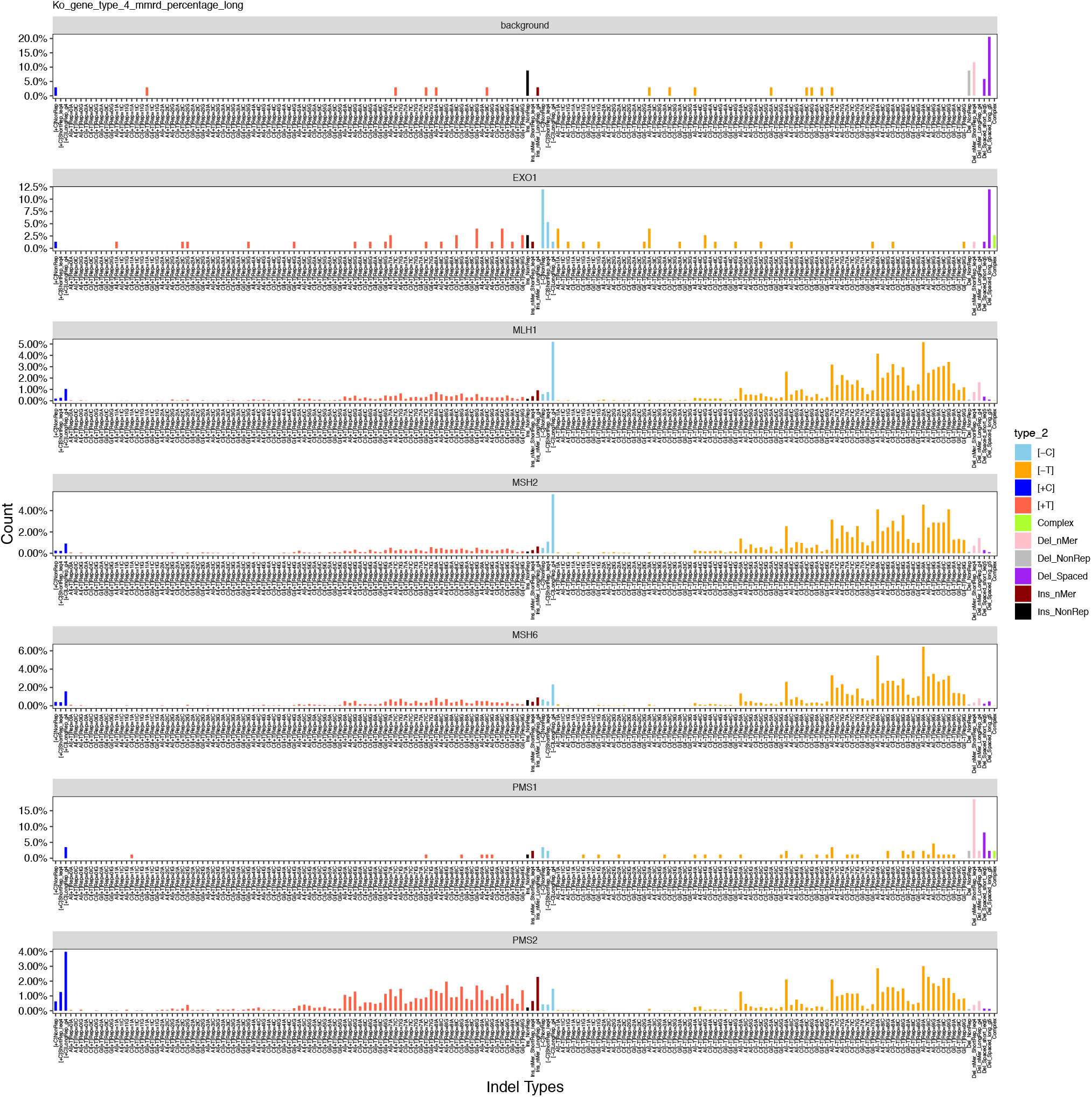
Indel signatures in 186 channels.

**Figure S9.**
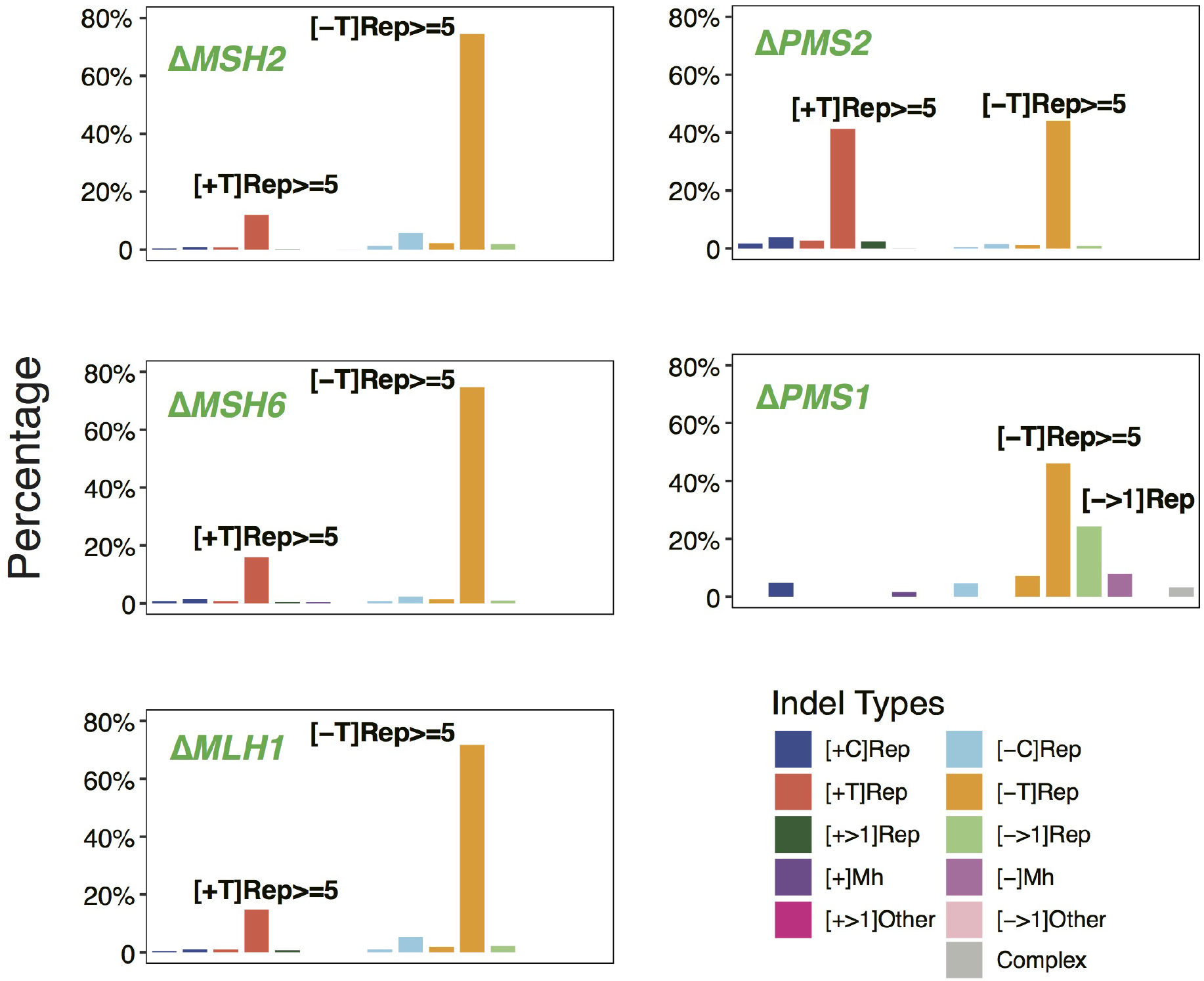
Indel signature of MMR gene knockouts in 15 channels.

**Figure S10.**
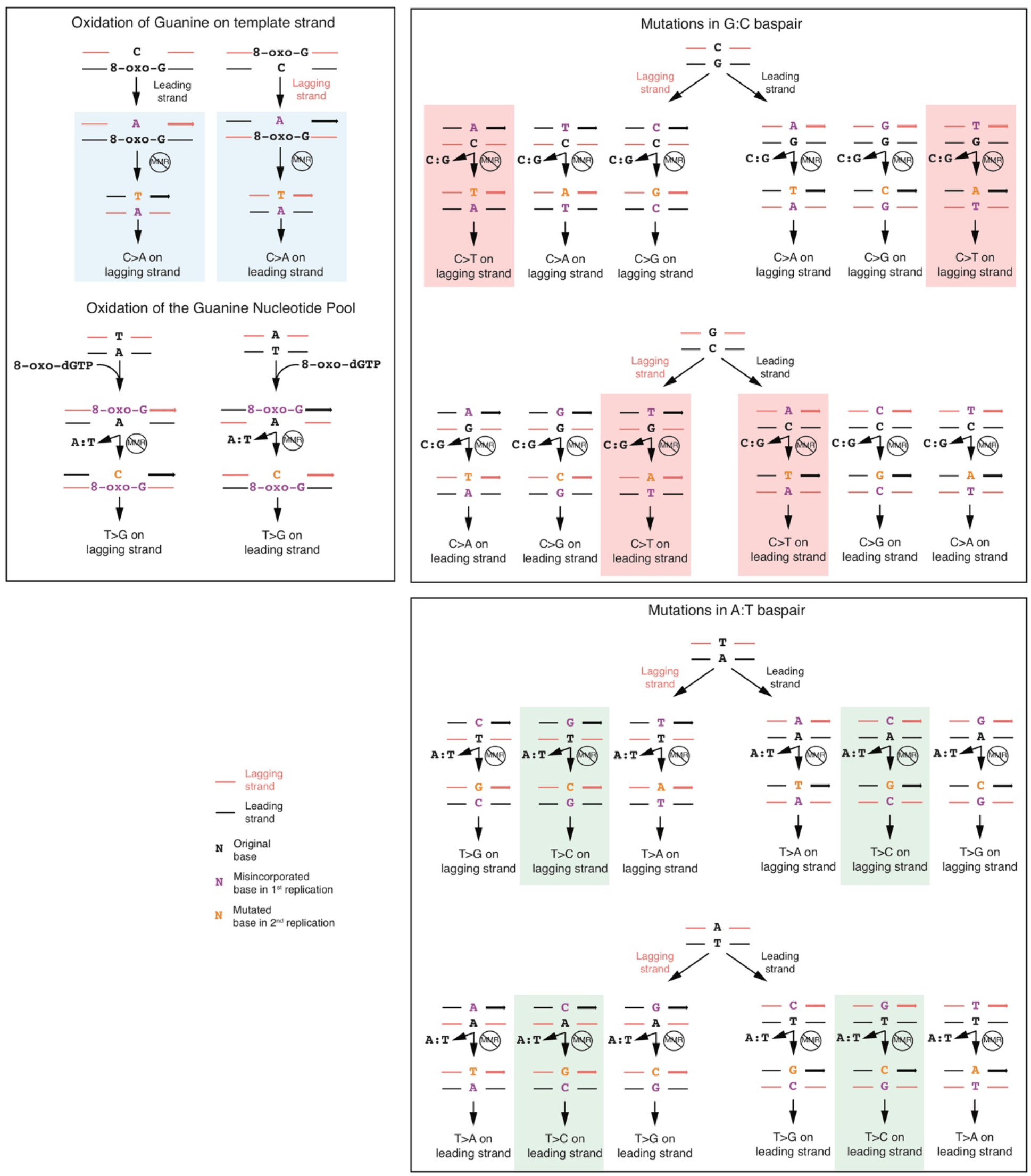
Putative outcomes of all possible base-base mismatches. Outcomes from 12 possible base-base mismatches. The red and black strands represent lagging and leading strands, respectively. The arrowed strand is the nascent strand. The highlighted pathways are the ones that generate C>A (blue), C>T (red) and T>C mutations (green) in the Δ*MSH2* mutational signature.

**Figure S11.**
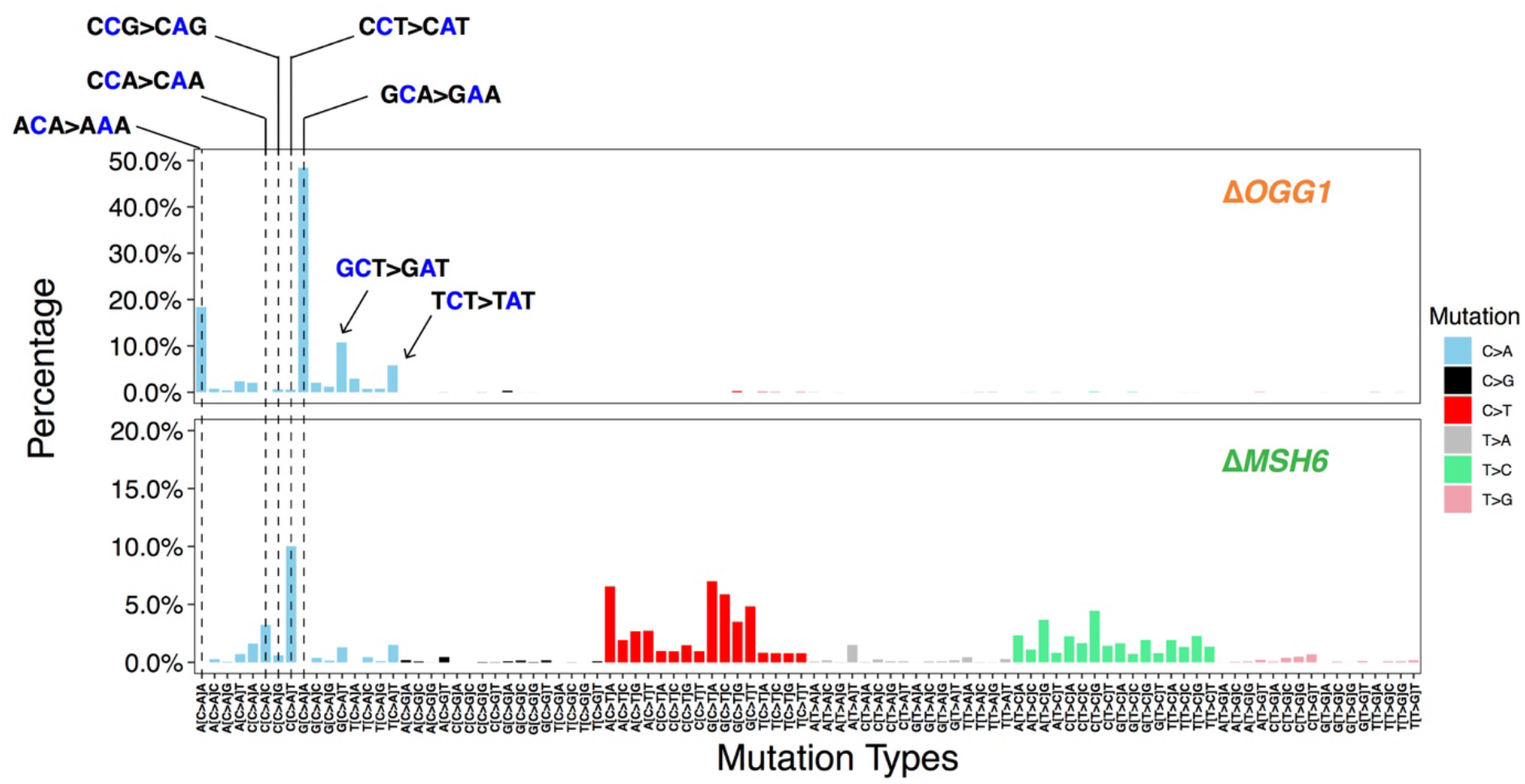
Comparison of trinucleotide context of C>A mutations generated by Δ*OGG1* and Δ*MSH6*.

**Figure S12.**
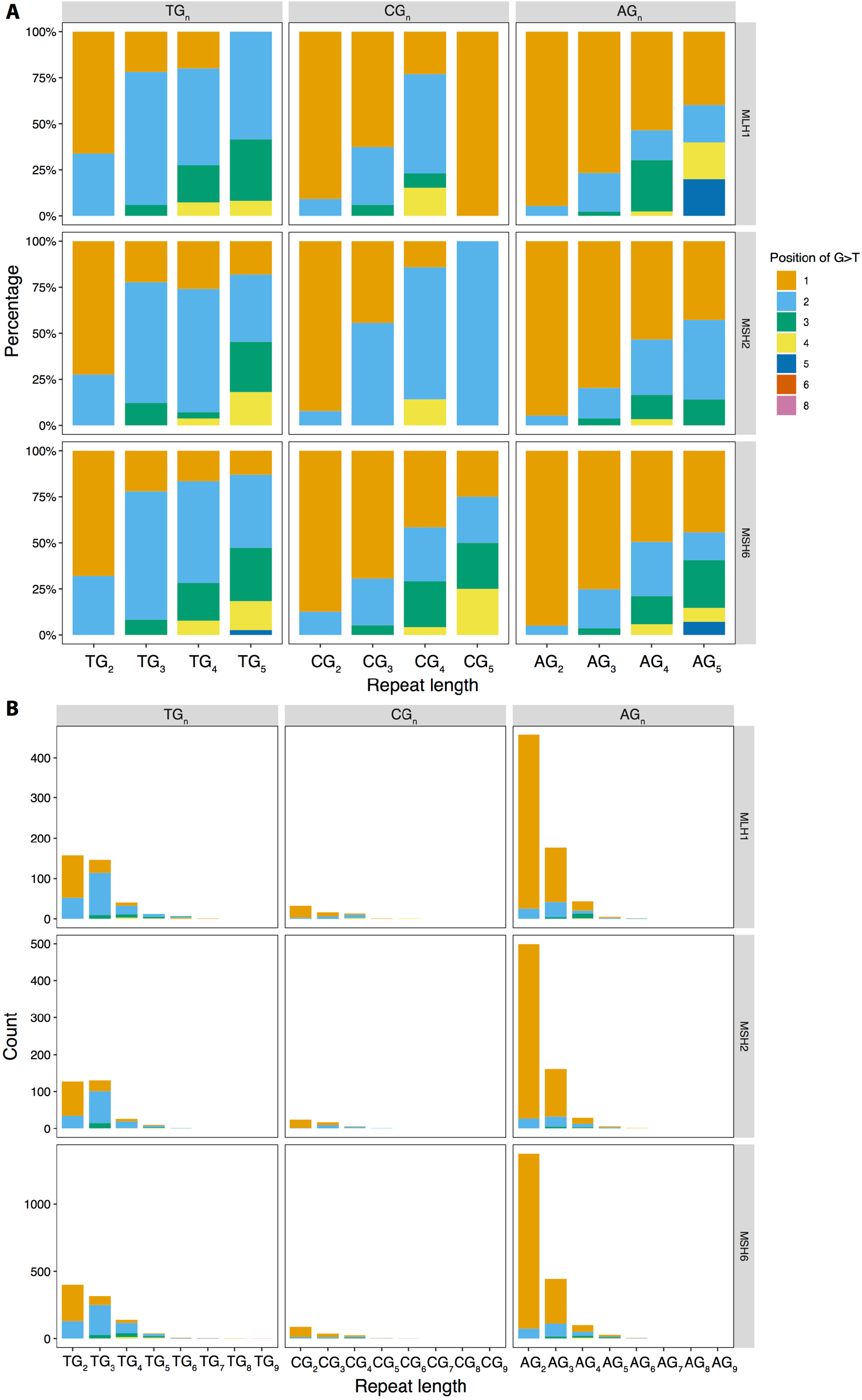
Distribution of G>T/C>A mutations in polyG tracts of MSH2, MSH6 and MLH1. (A) Relative frequency of occurrence of G>T/C>A in polyG tracts for Δ*MSH2, ΔMSH6* and Δ*MLH1.* (B) Occurrence of G>T/C>A in polyG tracts for Δ*MSH2, ΔMSH6* and Δ*MLH1*.

**Figure S13.**
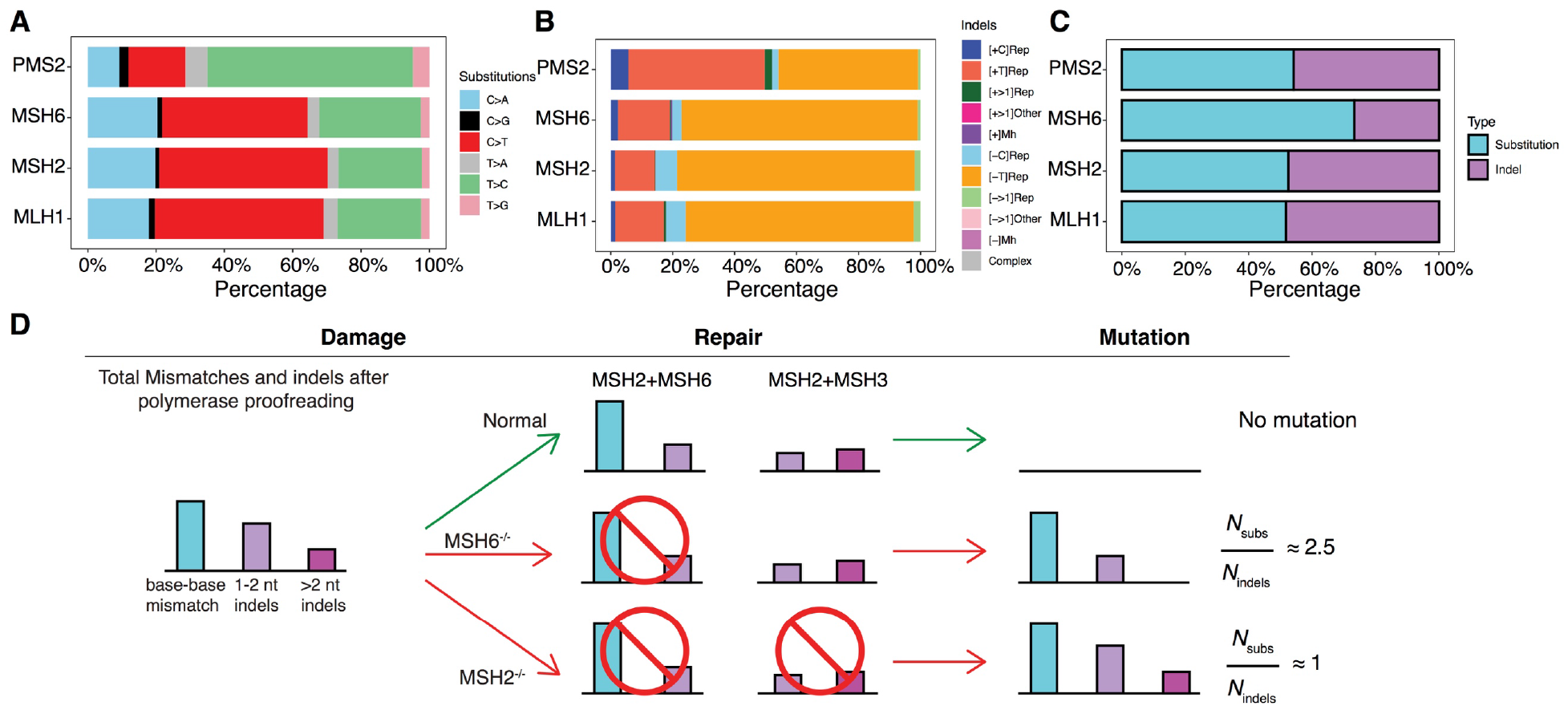
Proportion of different mutation types of substitution (A) and indel (B) signatures for 4 MMR gene knockouts. (C) The ratio of substitution and indel burden. (D) Schematic interpretation of the relative mutation burdens of Δ*MSH2* and Δ*MSH6*.

**Figure S14.**
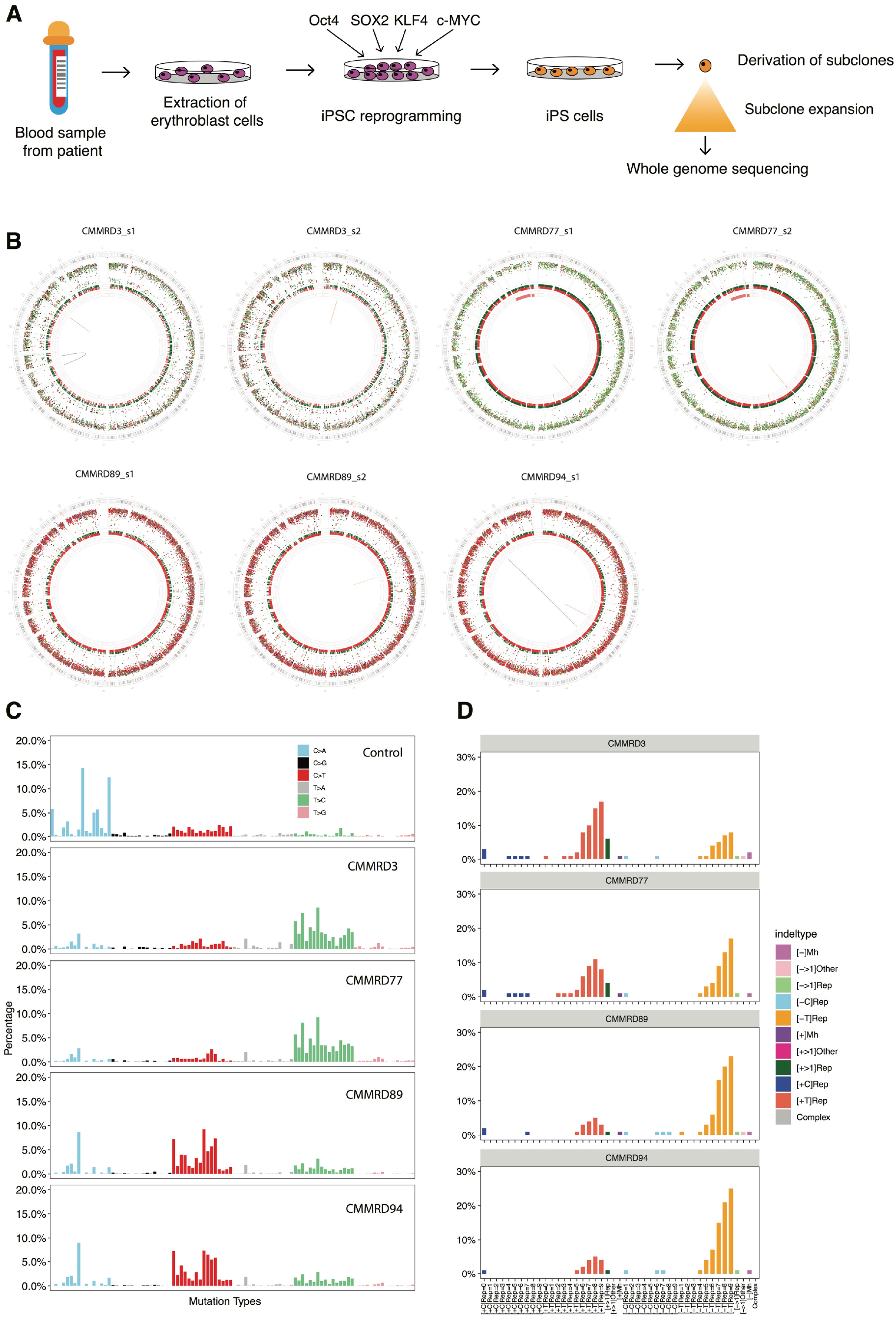
Mutational profiles of hIPSCs derived from patients with Constitutional MisMatch Repair Deficiency (CMMRD). (A) Experimental workflow used to generate hiPSCs from CMMRD patients, subcloning of hiPSCs and whole-genome sequencing. (B) Genome plots. Top: genome plots of four iPS cells from two PMS2 mutant patients. Bottom: genome plots of three iPS cells derived from two MSH6 mutant patients. Genome plots show somatic mutations including substitutions (outermost, dots represent six mutation types: C>A, blue; C>G, black; C>T, red; T>A, grey; T>C, green; T>G, pink), indels (the second outer circle, colour bars represent five types of indels: complex, grey; insertion, green; deletion other, red; repeat-mediated deletion, light red; microhomology-mediated deletion, dark red) and rearrangements (innermost, lines representing different types of rearrangements: tandem duplications, green; deletions, orange; inversions, blue; translocations, grey). (C) Substitution profiles. (D) Indel profiles.

**Figure S15.**
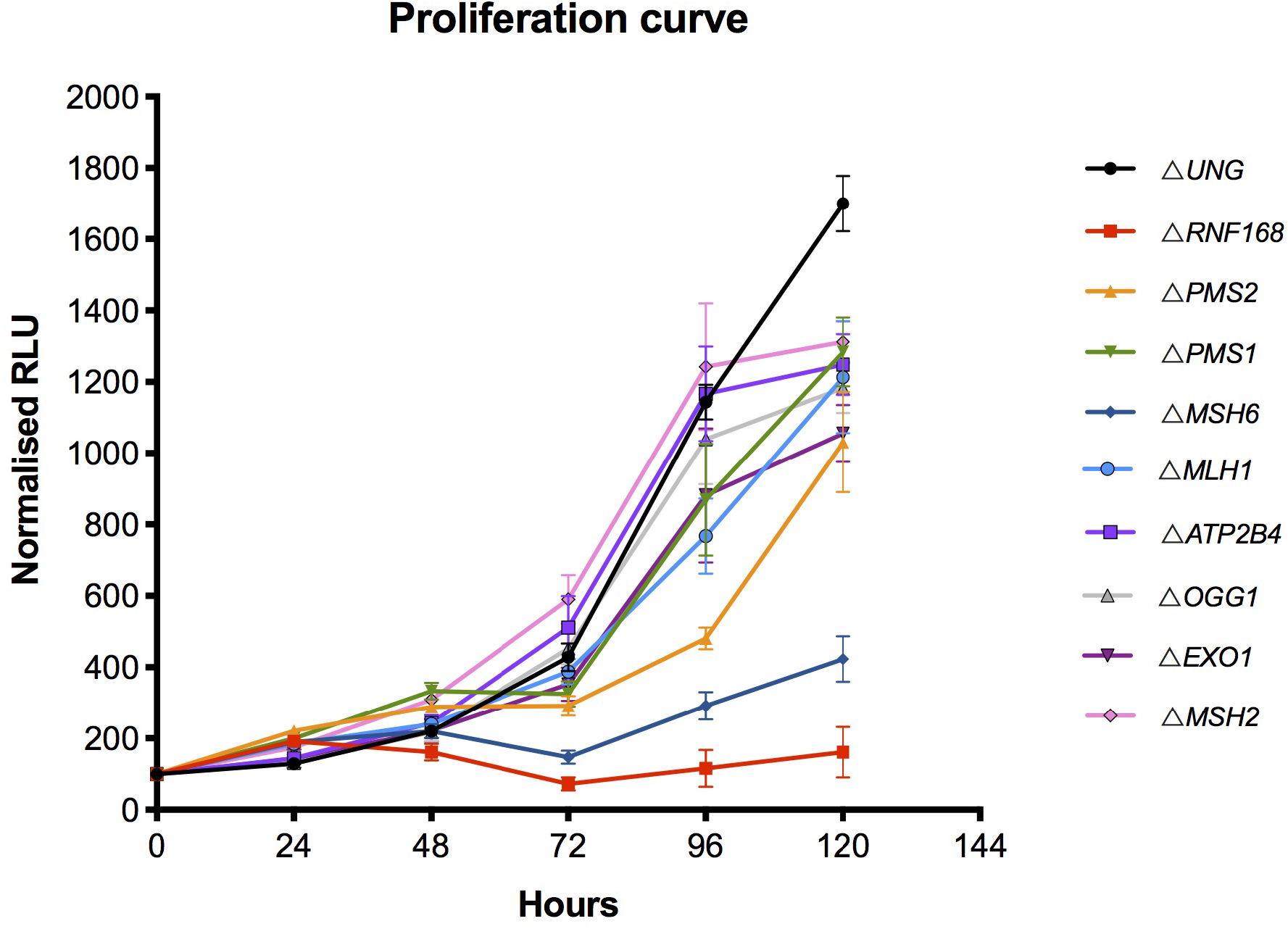
Proliferation of indicated knockout cell lines over a period of 5 days. Six replicates were used for each time point per indicated knockout lines. Error bars show standard error of the mean.

**Figure S16.**
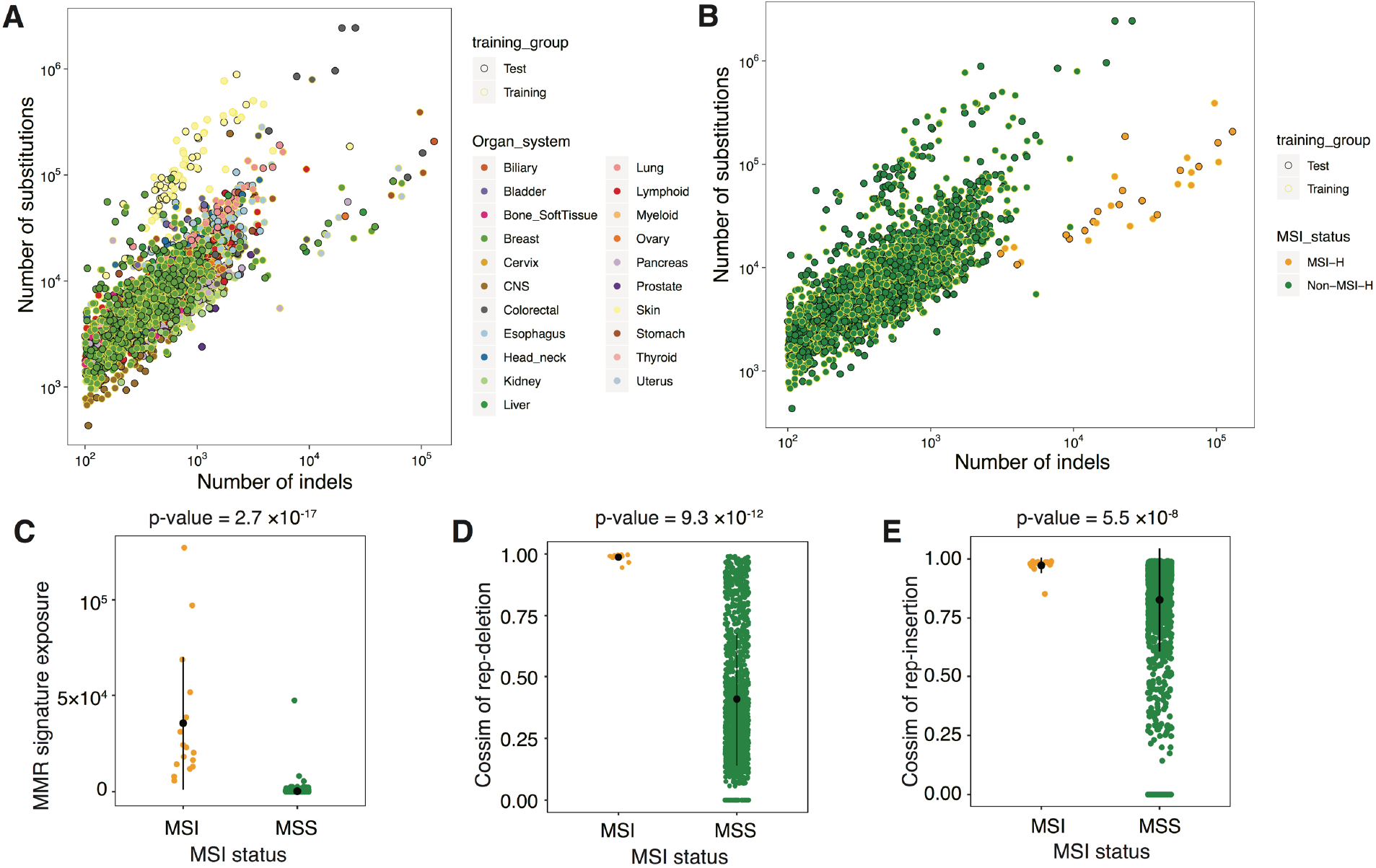
Parameters in MMRDetect.(A)(B) Distribution of mutation burden for 21 cancer types and MSI/non-MSI samples. Test and training data sets have similar distribution of mutations burden within different cancer types (A) and MSI status (B). (C)(D)(E) Distribution of the exposures of MMR-deficiency signatures and cosine similarities between repeat-mediated indels profiles of the tumor and MMR knockout indel signatures across MSI and MSS samples. Comparison of MSI and MSS on three features: (C) exposure of MMR signatures, (D) the cosine similarity between the profile of repeat-mediated deletions of cancer sample and that of knockout generated indel signatures, (E) the cosine similarity between the profile of repeat-mediated insertion of cancer sample and that of knockout generated indel signatures. P-values were calculated through Mann-Whitney test.

**Table S8.**
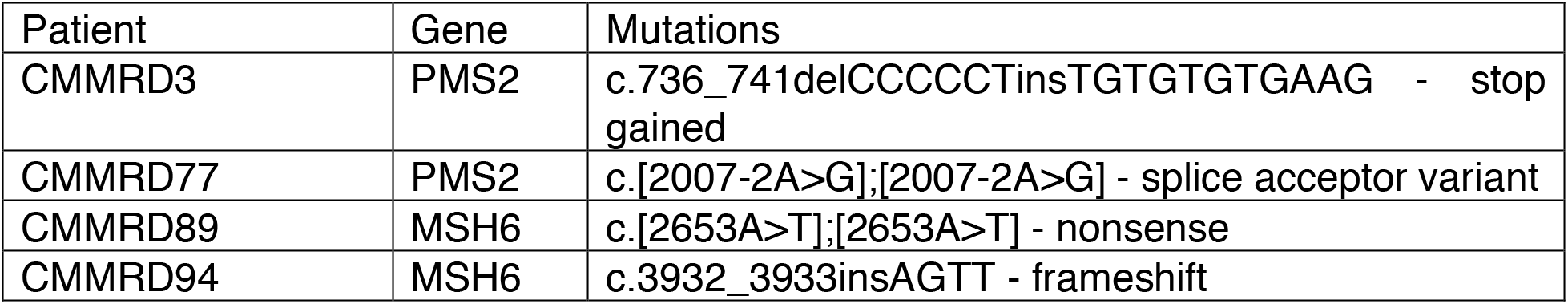
Genotypes of four CMMRD patients.

| Patient | Gene | Mutations |
| --- | --- | --- |
| CMMRD3 | PMS2 | c.736_741delCCCCCTinsTGTGTGTGAAG - stop gained |
| CMMRD77 | PMS2 | c.[2007-2A>G];[2007-2A>G] - splice acceptor variant |
| CMMRD89 | MSH6 | c.[2653A>T];[2653A>T] - nonsense |
| CMMRD94 | MSH6 | c.3932_3933insAGTT - frameshift |

## Supplementary Note 1

### Results of pilot study

A pilot experiment was performed to standardize the experimental procedure for the larger study. Three genes were selected for knockout (Δ): *MSH6* - a gene of the DNA mismatch repair (MMR) MutS family that produces strong substitution and indel patterns serving as a positive control, *UNG* – encodes for uracil-DNA glycosylase that eliminates uracil by cleaving the N-glycosylic bond and initiating base-excision repair (BER) which may or may not produce a mutational signature and *ATP2B4* - a gene for a primary ion transport ATPase playing a critical role in intracellular calcium homeostasis, unlikely to produce signatures and effectively a negative control. Two genotypes per gene were obtained and grown in culture to gauge reproducibility of signatures between different genotypes of a gene-knockout. These lines were cultured under normoxic (20%) and hypoxic (3%) states, for defined culture times of ~15, 30 or 45 days. Two single-cell subclones were derived for whole genome sequencing for each parental line (equivalent to four subclones per gene edit).

All parental clones and subclones were sequenced to > 30 fold depth. Short-read sequences were aligned to the human reference genome assembly GRCh37/hg19. All classes of somatic mutations were called in subclones subtracting on the primary iPSC parental clone. To ensure that mutational observations were not due to off-target edits, nor to acquisition of driver mutations, we searched for coding sequence mutations in parental clones and in subclones; we did not find any mutations of likely consequence. We also checked that intended edits had correctly targeted both alleles – all had achieved biallelic loss except one of the *UNG* genotypes appeared to be heterozygous, which was excluded in downstream analysis.

Comparing the *de novo* mutations accumulated in subclones after knocking out targeted genes, we observed detectable differences between these three gene knockouts *(ΔMSH6, ΔUNG* and Δ*ATP2B4*) in terms of mutation burden (Figure S1A) and mutation profile regardless of oxygen condition or length of time in culture. Furthermore, cosine similarity (cossim) between mutation profiles and parental clones (background) are clearly very dissimilar for Δ*MSH6* (cossim: 0.32-0.45) and Δ*UNG* (cossim: 0.67-0.88) again irrespective of oxygen conditions or time in culture and were identical for subclones derived from negative control Δ*ATP2B4* (Figure S1B). Similar observations were made for indels (Figure S2) where the burden and profile of mutagenesis was particularly detectable for the Δ*MSH6.*

Thus overall, the differences between normoxic and hypoxic conditions were not marked, although normoxic conditions produced slightly more mutations. Surprisingly, time in culture made only a marginal and non-linear difference to burden of mutagenesis (Figure S1A and S2A). Given the results of the pilot, weighing up the costs and risks associated with prolonged culture time (risk of infection, risk of selection, marked increase in cost of experimental reagents) with the minimal return in terms of mutation number, and also intending to minimize transitions between hypoxic to normoxic conditions while handling cell cultures, we opted to proceed with the fullscale study under normoxic conditions and for 15 days for the rest of study.

**Figure 1.**
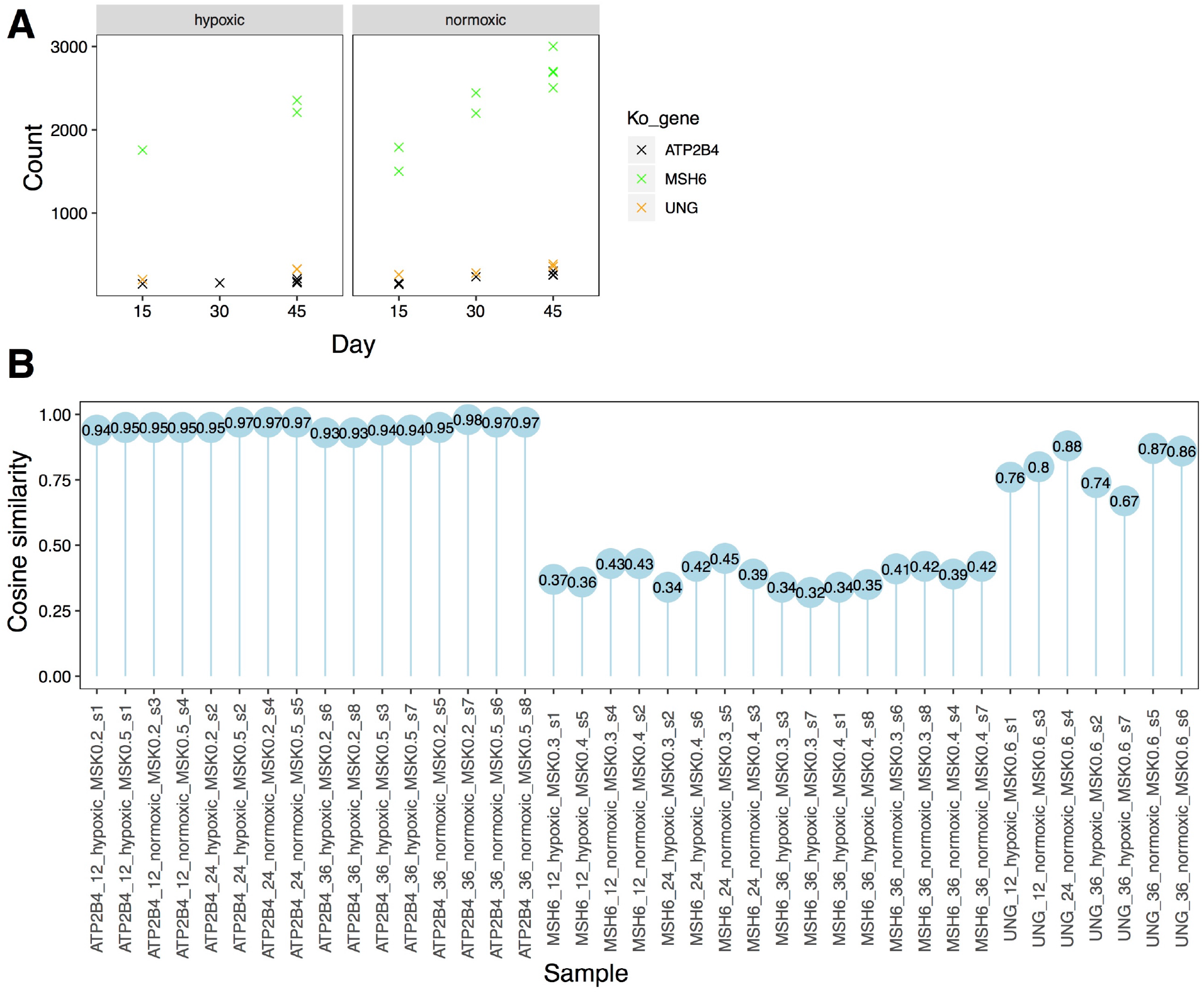
(A) Substitution burden for knockouts of ATP2B4, UNG and MSH6 under hypoxic and normoxic conditions as well as different culturing time. (B) The cosine similarities between the mutational profile of each subclone with background signature of culture.

**Figure 2.**
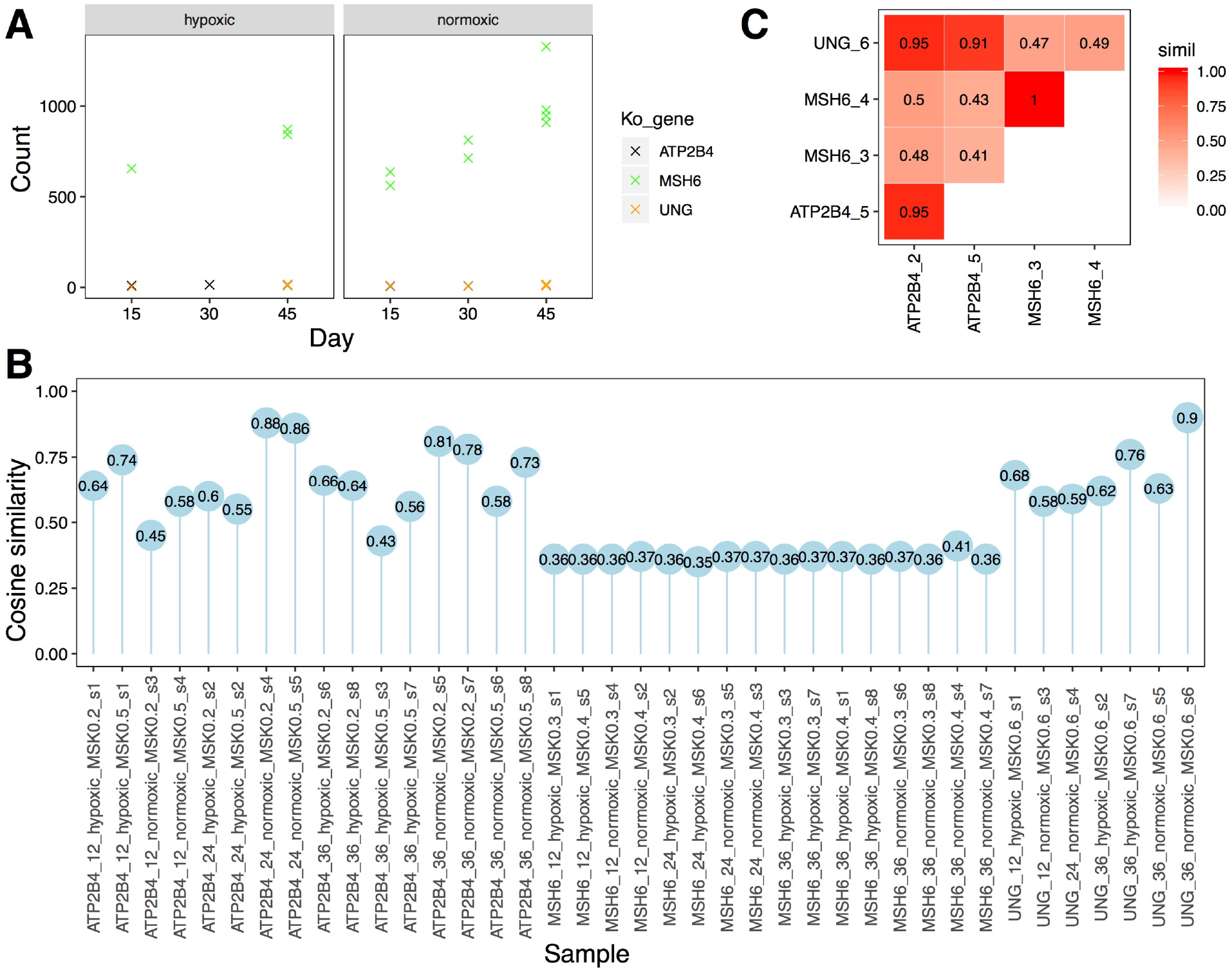
(A) Indel burden for knockouts of ATP2B4, UNG and MSH6 under hypoxic and normoxic conditions as well as different culturing time. (B) The cosine similarities between the mutational profile of each subclone with background signature of culture. (C) The cosine similarities between aggregated mutational profile of each knockout.

